# Safeguarding pollinators requires specific habitat prescriptions and substantially more land area than current policy suggests

**DOI:** 10.1101/2020.12.14.422715

**Authors:** Alana Pindar, Adam Hogg, Nigel E. Raine

## Abstract

Habitat loss and fragmentation are major drivers of global pollinator declines, yet even after recent unprecedented periods of anthropogenic land-use intensification the amount of habitat needed to support pollinators remains unknown. Here we use comprehensive datasets to determine the extent and amount of habitat needed. Safeguarding wild bee communities in a Canadian landscape requires 11.6-16.7% land-cover from a diverse range of habitats (~1.8-3.6x current policy guidelines), irrespective of whether conservation aims are enhancing species richness or abundance. Sensitive habitats, like tallgrass woodlands and wetlands, were important predictors of bee biodiversity. Conservation strategies that under-estimate the extent of habitat, spatial scale and specific habitat needs of functional guilds are unlikely to protect bee communities and the essential pollination services they provide to crops and wild plants.

**One sentence summary:** Safeguarding wild bee communities requires 11.6-16.7% of the area in common North American landscapes to provide targeted habitat prescriptions for different functional guilds over a variety of spatial scales.

## Main text

Human-induced land-use changes are driving unprecedented widespread and increasing global biodiversity losses (*1, 2*). These alarming declines in biodiversity result in the degradation of many essential ecosystem services and functions (*3, 4*), including pollination. Indeed, wild bees and the pollination services they provide to crops and wild plants are experiencing global declines in response to intensive anthropogenic landscape changes, climate change, parasites and diseases, competition from invasive species, and rising agrochemical usage (*5, 6*).

The Sustainable Development Agenda set globally agreed targets to end poverty, protect the planet, and ensure peace and prosperity for all by 2030 (*7*). However, less than a decade from this deadline little apparent progress has been made towards many of these key targets, including the need to ‘ensure the conservation, restoration and sustainable use of terrestrial and inland freshwater ecosystems and their services’ (Goal 15.1) (*7*). Efforts to slow, or even reverse global pollinator declines have led many countries to initiate conservation strategies in agricultural areas (*8–10*), urban environments (*11*), and other sensitive lands to mitigate the loss of vital pollinators and the ecosystem services they provide (*5, 12*).

Selection and implementation of specific conservation strategies will strongly depend on conservation priorities and may differ substantially if the goal is to: (1) enhance pollination by pollinators visiting particular crops (*13, 14*), (2) maintain wider pollinator biodiversity (*13)* or (3) specifically target the recovery of pollinator species-at-risk (*15*). Most research to date has focused on adding and restoring pollinator habitat, typically by planting more abundant and diverse floral mixtures as food sources (*16, 17*), and by providing or enhancing nesting sites and suitable larval host plants (*18*). Evidence suggests these strategies can be highly effective at increasing pollinator abundance and species richness (*8, 19*). While bee species richness and abundance are tightly associated with floral and nesting resources, these associations do not necessarily predict how much of a specific habitat is needed by any species (*19, 20*). Surprisingly, there is not yet any clear understanding of how much of each specific habitat type is required to support a healthy pollinator community, or indeed over what spatial scales such habitats are needed. This lack of information not only severely limits the ability to make and implement evidence-based recommendations to support pollinators at local or landscape scales, but also jeopardizes the chances of meeting globally agreed Sustainable Development Goals.

To address these critical knowledge gaps we compiled an extensive dataset of bees (~63,000 observations from 361 species, 86% of the species recorded from Ontario, Canada, from surveys over 15 years: 2000-2015) from sites with some degree of anthropogenic land use change (See Supplementary Information). In intensively managed and simplified ecosystems the provision of any additional suitable habitat will increase pollinator abundance and diversity (*19, 20*). However, at a certain point adding more habitat provides little or no further measurable pollinator biodiversity benefits (*21*)(Fig. 1b). We examined the shape of this relationship between the cumulative number of bee species supported when different amounts of suitable habitat are found in the landscape (closely following a species-area relationship) to find the point at which further additional habitat no longer enhanced species richness – a law of diminishing returns (Fig. 1b, Table S2). Unlike previous attempts at determining the relationships between bees and habitats within a landscape, our study used ground-truthed land cover data. This is critical as it provides greater confidence that habitat type designations from map datasets are meaningful descriptions of the reality of habitats (and critically the resources they provide to pollinators) on the ground. Furthermore, as different bee species can provide the same ecological function in different habitats (*13*), we determined both the extent of habitat and also the key habitat types required to maintain community species richness and abundance for five functional groups of bees (solitary ground nesters, social ground nesters, cavity nesters, bumblebees and cleptoparasites: see Supplementary Materials). Acknowledging that bee species can be highly mobile (*22, 23*) and require habitat resources at variable distances from their nests (*24*), we also tested which of 27 different habitat types were most widely used by bee communities at twelve different foraging ranges (in three categories: <500m, 750-1250m, >1500m) within a representative North American landscape (Fig 1a). To effectively demonstrate which habitats are most important to different functional groups of bees, and at what spatial scales, we mapped partial regression (β) coefficients of the most extensively used habitat types (reported in GLMMs) to generate habitat prescriptions that can be easily translated by all end-users into immediate best management practices and real-world conservation actions.

**Fig 1.**
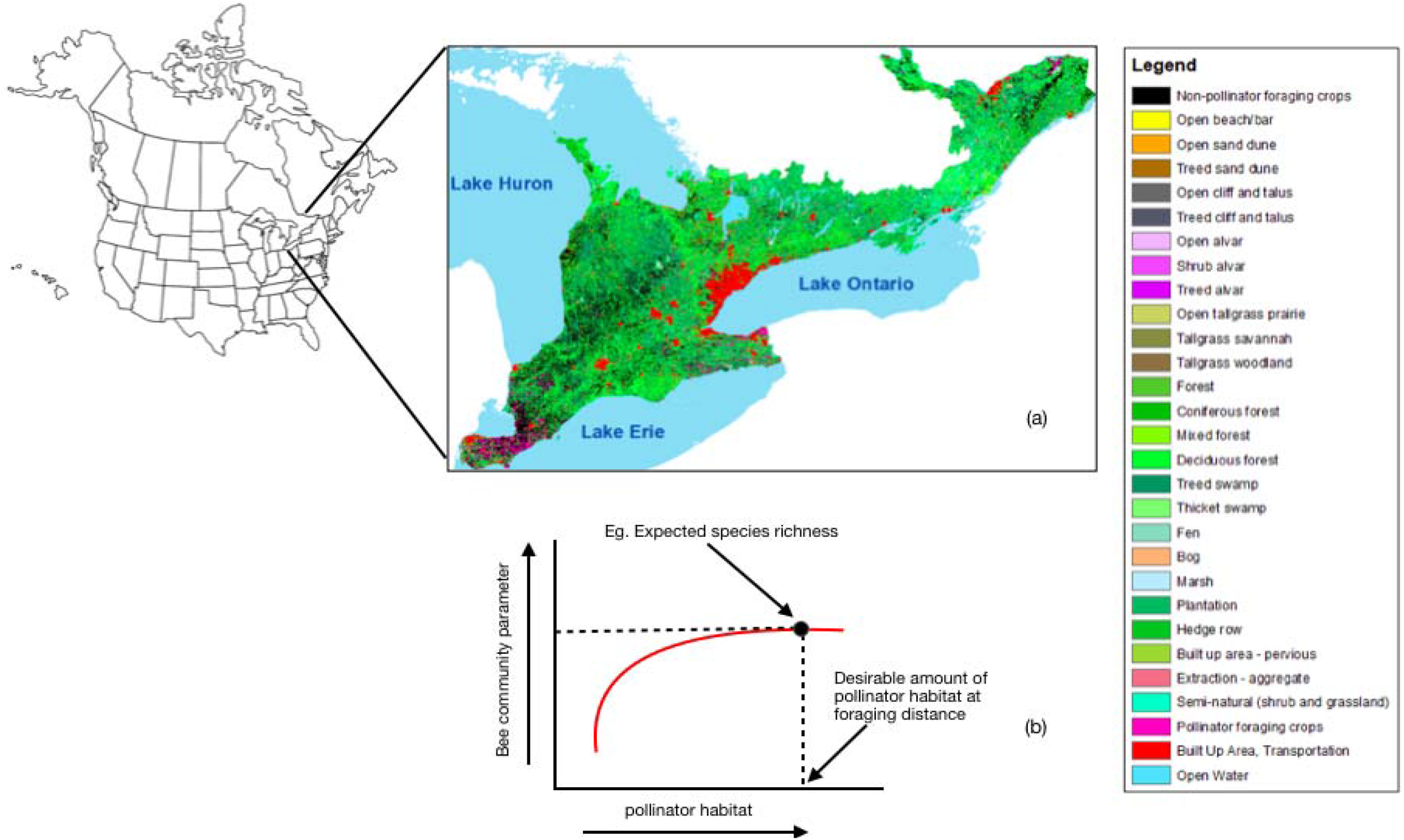
(a) Landscape gradient across Southern Ontario, Canada a typical North American landscape. Red (urban areas), black (intensive wind pollinated crops), and light blue (open water areas) reflect areas that provide little or no pollinator habitat. Pink represents intensive agricultural crops that provide pollinator foraging, while light- to darker-green colours represent a gradient of natural and semi-natural habitats. (b) The expected relationship between extent of pollinator habitat and the bee species richness supported in the landscape. Initial increases in the amount of pollinator habitat in a landscape are associated with a steep increase in bee species richness. However, the slope of this red line become less steep with additional increases in extent of pollinator habitat, until it reaches asymptote – the optimal landscape composition to support maximal bee species richness (marked with black dotted lines).

Our results suggest that conservation strategies to support wider bee biodiversity should preserve 11.6-16.7% of the land area as suitable habitats within a North American landscape (Fig. 2; 750-1250m). Current policy recommendations suggest maintaining 6% habitat in UK farmland to support pollinators based on the expected resource requirements for six common crop-visiting bee species (*9*) and an aspirational target to conserve 4.5% of habitat to support pollinators in Ontario, Canada. Compared to our results, both these policies substantially under-estimate (by 1.8-3.6 times) the amount of habitat needed to support diverse bee communities and safeguard the pollination services they provide. Any strategies aiming to safeguard pollinator biodiversity using targets below our evidence-based recommendations will likely provide insufficient habitat for wild bees.

**Fig 2.**
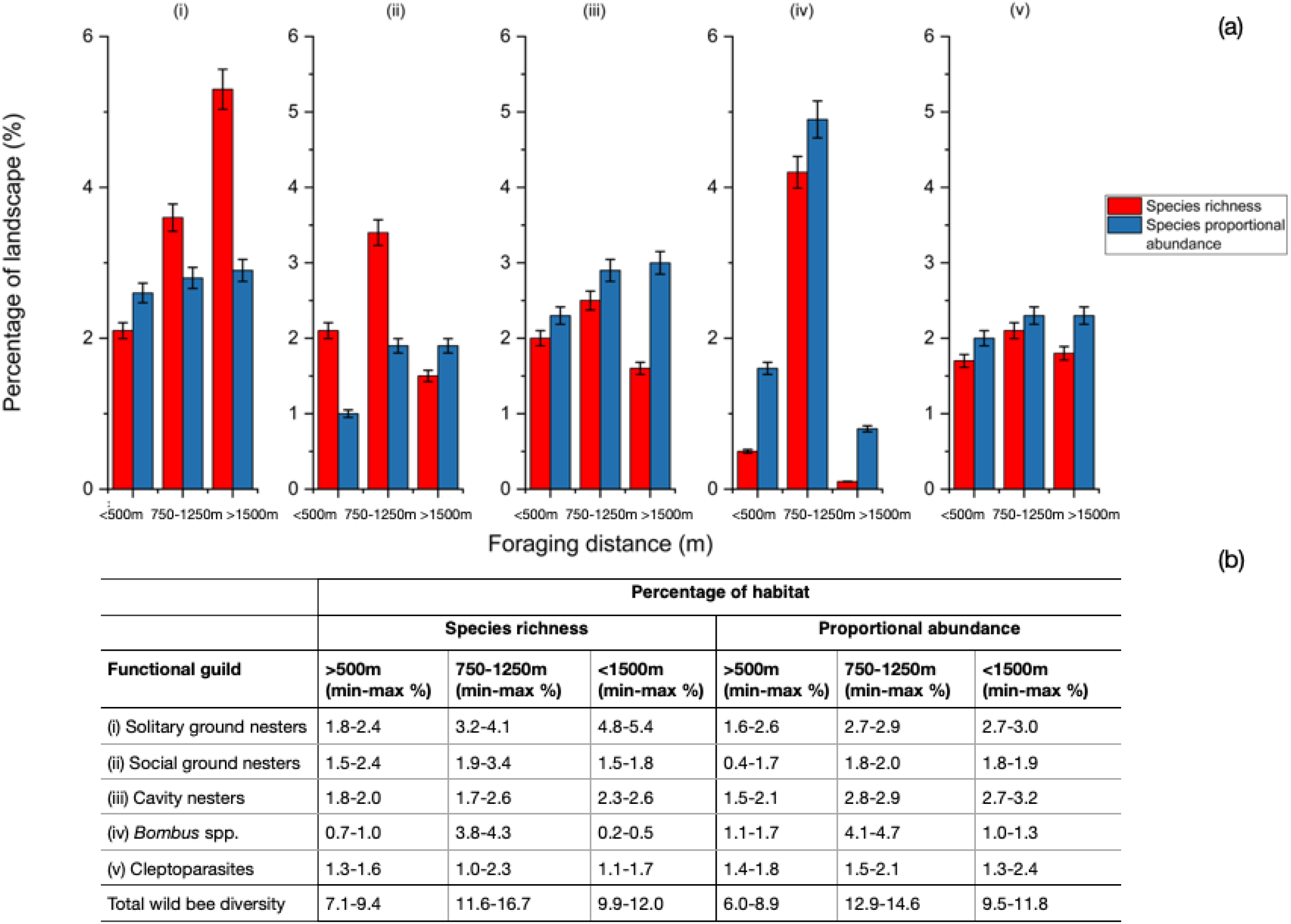
Extent of habitats required within a landscape to maintain the species richness (red-) and proportional abundance (blue columns) of five functional bee guilds: (i) solitary ground nesters, (ii) social ground nesters, (iii) cavity nesters, (iv) bumblebees (*Bombus* spp., and (v) cleptoparasitic species expected community parameters at each foraging category (<500m, between 750-1500m, and >1500m).

The full heat map clearly shows a diverse range of habitat types are needed to support wild bee communities (Fig. S3). However, to help end users successfully prioritise the most important habitats to maintain, restore or create we filtered the full heat map (by removing habitat types with interquartile ranges <0.25 for significant β coefficients) to highlight the most important habitat types in a landscape (Fig. 3). If the goal is to safeguard wider pollinator biodiversity, more habitat and distinctly different habitat types are required (Fig. 3; Fig. 4a-e) than if the goal is to enhance crop pollination through increasing the abundance of specific functional groups or indeed particularly important species (*25*) (Fig. 3; 4f-j). Specifically, more habitat types occupying an increased percentage of the landscape at larger spatial scales would need to be provided to support a greater richness of solitary ground-nesting species in comparison to if the goal is to maintain their community abundance composition (Fig. 2ai, 4a,f). Functional groups, other than cavity nesters and cleptoparasites, showed a preference for habitat at foraging distances between 750-1250m over more localized (<500m) and more dispersed scales (>1500m) (Fig. 2a). It is likely that the lack of observed changes in the amount of habitat needed to conserve cleptoparasitic species richness and abundance with respect to distance is because their distribution will be strongly influenced by the habitat preferences of their host species (Fig 2a, b) (*26*).

**Fig 3.**
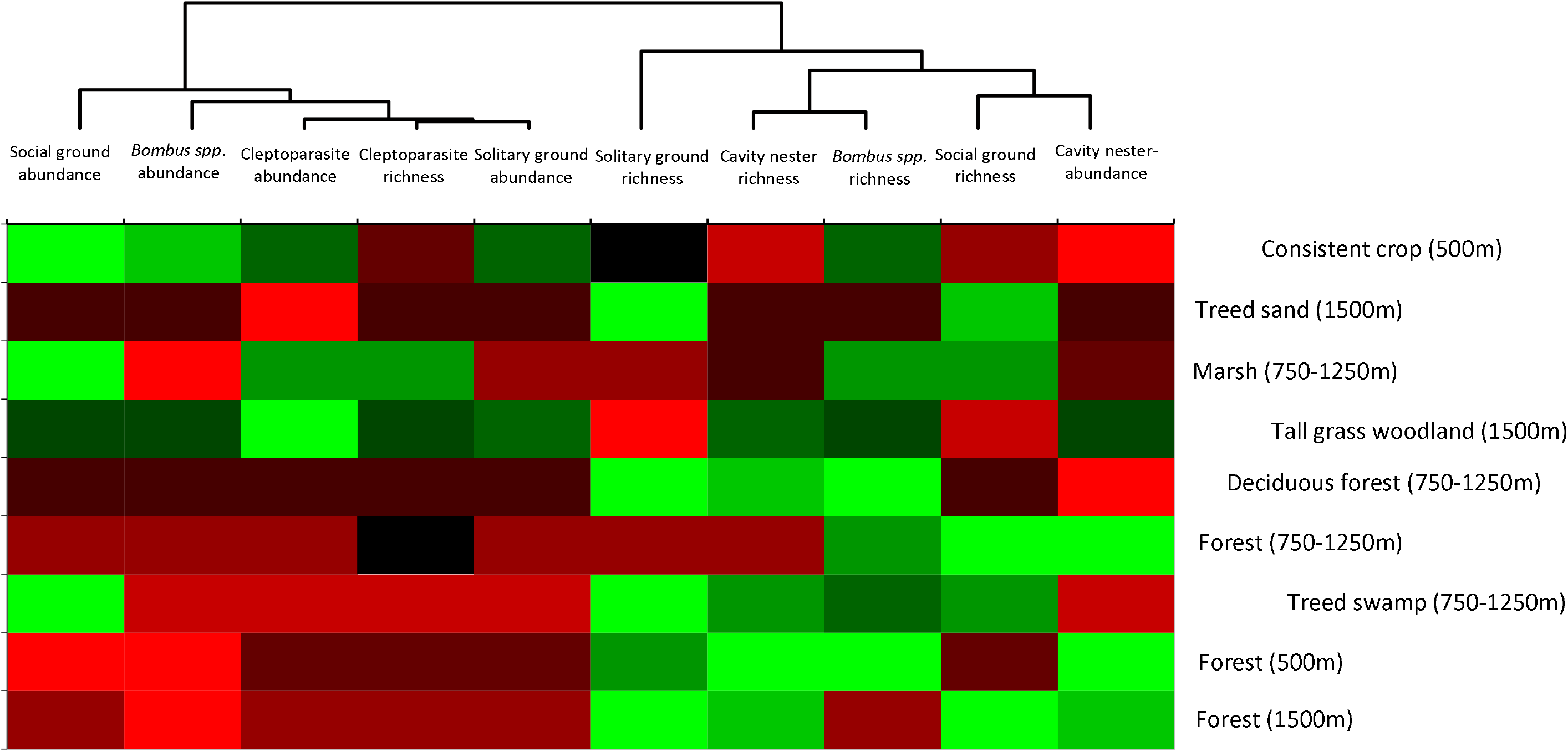
Heat map showing the most important habitat types driving key bee biodiversity metrics (species richness and proportional abundance) of five functional bee guilds: solitary ground nesters, social ground nesters, cavity nesters, bumblebees (*Bombus* spp.), and cleptoparasitic species at three foraging distance categories (<500m, 750-1500m, and >1500m). Lighter shades of green indicate greater use of the habitat at different spatial distances, where darker shades of red suggest a less desirable habitat for supporting functional guild species richness and/or abundance. Habitat similarity is characterized by groupings of alike colours, either among function guilds (horizontal rows) or across spatial distances and habitat types (vertical columns). Forested habitats represented 50m of habitat edges. This is a filtered version of the overall heat map (Fig. S3) from which habitat types with an interquartile ranges of <0.25 of significant β coefficients (habitat types) have been removed. Black cells indicate the habitat has a neutral impact on bee species richness and abundance in the landscape.

**Fig 4.**
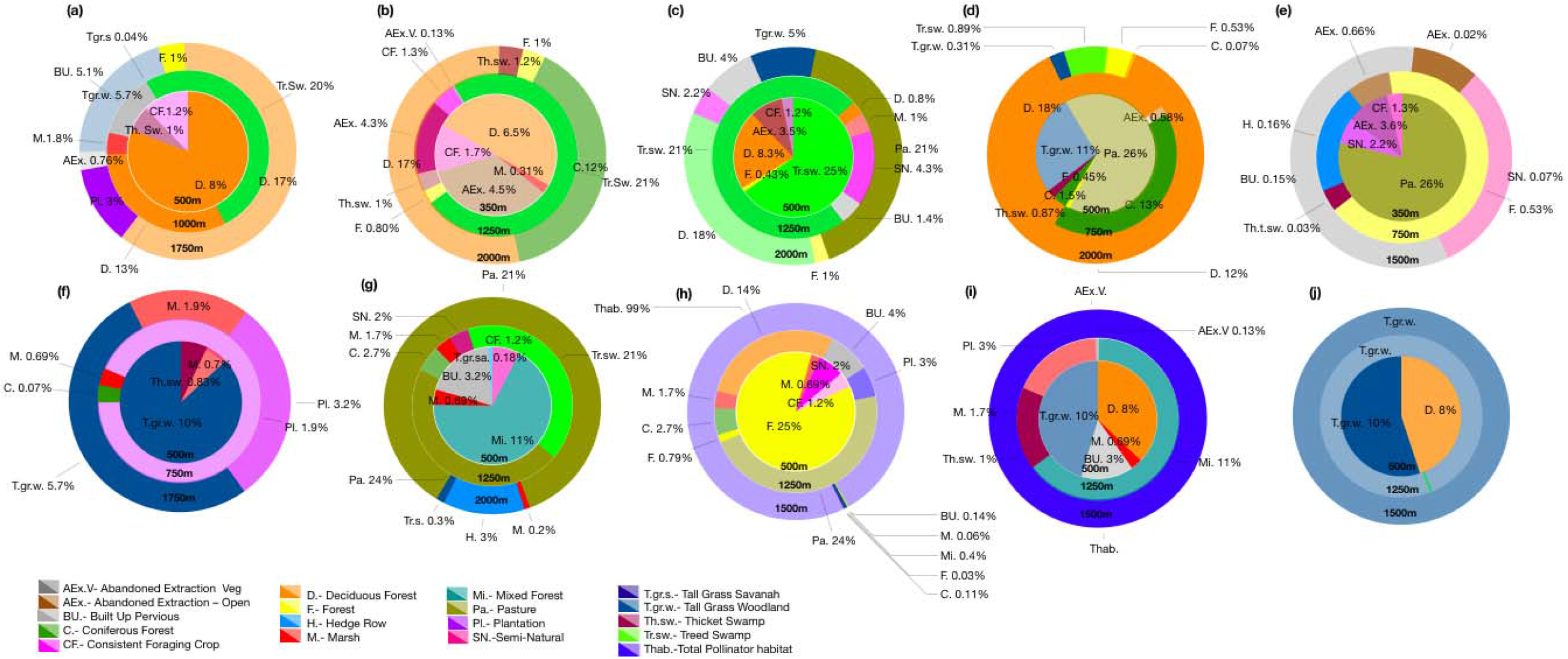
The most preferred habitat types and required percentages at each spatial scale within spatial categories (<500m, 750-1250m, and >1500m) for (a-e) species richness and (f-j) abundance of five bee functional guilds: (a, f) solitary ground nesters; (b, g) social ground nesters; (c, h) cavity nesters; (d, i) bumblebees (*Bombus* spp.); and (e, j) cleptoparasites. Colour shades among habitat types illustrate significant coefficients reported in Tables S8-10. Lighter shades represent significant negative coefficients in models, which represents less critical, but not unpreferred, habitat types.

Many of the identified pollinator species-at-risk in North America are bumblebees (*27*). Given that these major crop pollinators showed considerable preferences for habitat between 750-1250m in our study (250-1000m in the UK: *28*), we suggest that implementing agri-environmental conservation schemes in North American landscapes that focus on ensuring natural/agricultural pollination resources at habitat distances of <750m will likely miss opportunities to enhance pollination services provided by wild *Bombus* species (Fig. 2a, b). The importance of conserving sensitive lands, such as tallgrass woodlands and wetland habitat, for bee species appeared to far outweigh other habitat types such as hedgerows and semi-natural habitat (Fig. 3; Fig 4). Wetland and forest edge habitats were significant predictors of species richness in all bee groups across a range of foraging distances (Fig. 3; Fig. 4).

Promoting and maintaining a variety of forest edge habitats in agricultural areas where *Bombus* species and cavity nesters are the predominant crop pollinators could represent a more effective strategy to increase crop pollination services than implementing flowering field margins that may provide less varied nesting opportunities for these target groups (Fig. 3, Fig. 4b, d, g, i). Given that many habitat losses in North America are the result of natural land conversion to agricultural uses (*29, 30*), and that increases in agricultural habitat in landscapes have resulted in significant loss of phylogenetic diversity in bee communities (*31*), it is important that environmental policy in agricultural landscapes consider addition, restoration or creation of wetland habitats. The best conservation policies for supporting pollinators may also deliver other biodiversity benefits, for example providing suitable habitat for other beneficial arthropods (e.g. spiders and parasitoid wasps that can provide crop pest bio-control), birds and other wildlife in the landscape. The ecosystem services provided by wetlands extend far beyond pollinators - wetlands increase water table height and therefore quantity of water available for crop irrigation, improve drinking water quality, flood mitigation and provision of habitat for other wildlife, including other species-at-risk (*32*).

It is critical to continue to implement wild pollinator monitoring programs and to identify specific ecological requirements for individual pollinator species before and after the implementation of conservation strategies. Such monitoring programs will be the best indicators of how populations are responding to any new or modified management practices. Overall, we still know very little about the foraging patterns and flower preferences of the majority of wild bee species (*33*), although some species (e.g., *Eucera* (*Peponapis*) *pruinosa*, *Nomia melanderi*, and common bumble bee species) are comparatively well studied.

In the face of evidence that intensive landscape management can severely limit the diversity and extent of habitat to support wild pollinators (*3,5*), global conservation policies must not under-estimate what the pollinators actually need to survive and thrive. Our results provide clear-cut habitat prescriptions to support specific conservation needs for wild bees. As a society we need to have a clear understanding of the specific aims, priorities and outcomes required for pollinator conservation with regards to crop pollination, maintaining wider biodiversity or targeting key species-at-risk. Our results clearly highlight that whether supporting species richness or abundance, the wrong habitat prescription will ultimately continue to prove ineffective for safeguarding wild pollinator biodiversity and the essential ecosystem services they provide.

## Acknowledgements

We would like to thank Mace Vaughan and Laurence Packer for constructive comments and thoughtful discussions on earlier versions of this manuscript. This work was funded by the Natural Sciences and Engineering Research Council of Canada (NSERC) Discovery Grant (2015-06783) and the Food from Thought: Agricultural Systems for a Healthy Planet Initiative, by the Canada First Research Excellence Fund (grant 000054). A.P. was supported by the Webster Postdoctoral Fellowship in Environmental Sciences, University of Guelph and by the Weston Family Foundation. N. E. R. was supported as the Rebanks Family Chair in Pollinator Conservation by the Weston Family Foundation. We dedicate this manuscript to the memory of Gordon Taylor, Alana’s Dad and constant source of inspiration.

## Author Contributions

A.P., A.H., and N.E.R designed the research and analyzed data. A.P and N.E.R wrote the paper.

