## Supplemental Materials for "Safeguarding pollinators requires specific habitat prescriptions and substantially more land area than current policy suggests"

Alana Pindar^1†^, Adam Hogg^2^, Nigel E. Raine^1^

^1^ School of Environmental Sciences, University of Guelph, Guelph ON, N1G 2W1, Canada

^2^ Ministry of Natural Resources and Forestry, Peterborough ON, K9J 8M5, Canada

**1.0 Creating a wild bee database for Ontario**

Recent estimates of bee diversity suggest that the province of Ontario is home to 421 of the 926 bee species found in Canada, making Ontario a national bee biodiversity hotspot and a critical location for strong pollinator conservation policy (*33-35*). Ontario is one of the few provinces or territories in Canada in which multiple survey studies of wild bees have been conducted. We compiled a database that includes data from 34 wild bee surveys conducted in Ontario between 2000-2015 and consists of 66,271 individual bee specimen records, representing 34 of the 39 genera, and 361 species out of a possible 421 species (85.7% of the known species) recorded for the province (Table S1). Our database includes no species records from five other genera known from other records in the province: *Svastra* spp. (Apidae); *Pthilothrix bombiformis* (Cresson) (Hymenoptera: Apidae); *Dieumonia* spp. (Halictidae); *Dufourea* spp. (Halictidae); and *Dianthidium* spp. (Megachilidae) (Table S1)(*34, 36*). All bees reported in each study/survey were included in the database with the exception of a small number of *Nomada* specimens. These specimens were flagged and removed from the database due to their unreliable or unverified identification information (for example, “*Nomada* spp*./* spp1/ Form a/ Form d/ Form e”) to avoid confusion or duplication of individual or species records.

*1.1 Quantifying bee community biodiversity targets in a common North American landscape*

We chose to use bee species richness (number of species present) and abundance (number of individuals present) as target parameters for this study as both are widely used fundamental metrics of assessing changes in biodiversity across time and space (*37*). An additional benefit to using simple biodiversity metrics, like the numbers of individual bees present and the number of species recorded (rather than more derived biodiversity parameters like species diversity and evenness) is that this ensures our results are more understandable and directly relevant for policymakers, farmers, land owners, conservationists and other important stakeholders that will use this information to implement changes to promote pollinator conservation on the ground.

All measures of species richness and individual abundance in our analyses exclude honey bees (*Apis mellifera*) because any of these caught in surveys were almost certainly from managed colonies. To account for differences in sampling effort and technique among the 34 studies/surveys included in the database, we assessed whether our bee biodiversity parameters (species richness and abundance) reported in each survey differed from random. To do this, we created an expected distribution for each survey by randomly sub-sampling the same number of individuals collected in a survey from the full database of 66,271 individual specimen records. All individuals were equally weighted to allow the same opportunity of sampling individuals with different social structures and nesting preferences.

While functional diversity is technically defined as the diversity of species’ traits, this term is also often used to represent the diversity of species’ ecological functions and/or guilds within an ecosystem (*38, 39*). Unlike traditional metrics used to measure species richness and diversity, metrics of functional diversity provide a mechanistic link between species and environmental factors (*39*) as species are assigned to a guild based on *a priori* expectations of taxonomic, morphological, or physiological traits (*40*). Therefore, using functional guilds to assess the status of bees in diverse habitats in Ontario provided us greater insight into the ecological habitat requirements of different bee species compared to examining bee communities as a whole (*41, 42*). Re-sampled bee lists from each of the 34 studies were divided into the following guilds for analysis: (1) solitary ground nesters, (2) social ground nesters, (3) cavity nesters, (4) bumblebees or *Bombus* spp. (except subgenus *Psithyrus*), and (5) clepto- and social parasites (including *Bombus* subgenus *Psithyrus*). Ground nesters were split into two guilds because social bee colonies can contain many individuals per nest and are active over a much longer period of the year, whereas most solitary bees have only one female per nest and are generally active for only a few weeks (*42, 43*). **Solitary ground nesters** (including species in the genera *Agapostemon*, *Andrena,* *Colletes, Lasioglossum* subgenus *Dialictus* [other than the species listed under cavity nesters below; although many *Dialictus* are eusocial, those we found in this study are solitary (*44*)], and *Lasioglossum* subgenus *Lasioglossum*) and **Social ground nesters** (including species in the genera *Augochlorella*, *Halictus*, and *Lasioglossum* subgenus *Evylaeus*) generally prefer open habitats, often those with dry sandy soils. **Cavity nesters** (including species in the genera *Augochlora,* *Ceratina*, *Hoplitis*, *Hylaeus, Lasioglossum (D.) cressonii* Robertson*, Lasioglossum (D.) semicaeruleum* Cockerell, and *Osmia*,) nest in pithy plant stems, rock cavities and abandoned beetle burrows in wood, usually using pre-existing cavities (*45*). ***Bombus* spp.** (bumblebees) were placed in a guild on their own, as they are social cavity nesters. Because species within *Bombus* subgenus *Psithyrus* are social parasites, the *Bombus* guild refers only to the non-parasitic species. **Cleptoparasites** (species in the genera *Nomada* and *Sphecodes*) as well as social parasites (*Bombus* subgenus *Psithyrus* spp.) were united under the guild “cleptoparasites” as they are all bees that lay their eggs in nests of other bee species.

We tested for significant differences between expected and observed species richness and abundance for each of the 34 surveys/studies using a nonparametric median test (i.e. mood test). Despite high variation of observed bee species richness and abundance among survey sites and over time, these differences between observed and expected values were only significant for solitary, social ground nester species, and cleptoparasite species richness (Fig S1a; Table S2). There were no significant differences between observed and expected species richness for cavity nesters or *Bombus* spp. (Fig. S1a) nor proportional abundance for any of the functional guilds (Fig. S1b; Table S2). These results were robust to variation in the number of times we randomly sampled species from the overall dataset of 66,271 observations (Table S2), suggesting functional roles within a pollinator community remain largely consistent across a landscape although the particular species found within specific habitat types vary (*37*). We suggest that any strong deviations from our reported observations in other studies likely indicate that bee communities are suffering from environmental perturbances or year-to-year variability.

**2.0 Quantifying pollinator habitat needs**

*2.1 How much habitat is needed?*

Although some studies have shown the importance of specific habitat types, such as semi-natural and natural habitat, urban, and consistent foraging crops, for particular bee species at various spatial scales in a landscape (*46-51*) very little information currently exists around how much pollinator habitat is actually required to maintain species richness or abundance for bee communities. Here we define pollinator habitat as land that meets all of the forage, nesting and hibernation requirements of wild bee species (*52*). Adding pollinator habitat, such as wildflowers and nesting resources, to a landscape can be achieved in a variety of ways. For example, recommendations from the UK suggest planting 1-4 hectares of flowering forage and providing 0.5-2 hectares of nesting resources per 100 hectares of farmland is enough to provide six common pollinator species (including three solitary (*Andrena* spp.) mining bee and three bumblebee species) with sufficient nesting and foraging opportunities to thrive (*49*).

In 2016, the government of Ontario, Canada introduced policy as part of the provincial Pollinator Health Action Plan to “restore, enhance and protect, one million acres of pollinator habitat” (*53*), which represents about 4.5% of the land area of Southern Ontario (mixed wood plain ecozone) where most of province’s rich agricultural lands are situated. Whilst one million acres (or 404,686 hectares) sounds like an impressively large area, there is no evidence to suggest that this much pollinator-suitable habitat will be sufficient to conserve healthy communities of wild pollinators and the essential ecosystem services they provide for agricultural production (*54-56*) and native plant communities (*57*) in a landscape. Using a comprehensive land use data set from Ontario, representing 24 habitat types (Tables S3, S4) at three different spatial scales (<500m, 750-1250m, >1500m), we examined whether this government policy is adequate to safeguard the abundance and richness of bee species.

To determine the total amount of habitat needed to maintain species richness and abundance (target bee community parameters) for each functional guild (solitary ground nesters, social ground nesters, cavity nesters, *Bombus* spp. and cleptoparasites), we determined the proportion of pollinator habitat found from the overall area at each foraging distance (the radius of the circle centred on the study site location) in relation to the target bee community parameter calculated from resampling the 34 bee surveys (Fig. S2). The mean amount of habitat needed to maintain bee community parameters for each functional guild per spatial category was analyzed using equation 1:

*ln(x_i_)= E(a, r)- β*_0_	*(Equation 1)*

*β*_1_

where *E(a, r)* is the expected proportional abundance (*a*) and species richness (*r*) of functional guilds (solitary ground, social ground, cavity nesters, *Bombus* spp*.* and cleptoparasites), *β*_0_ is the slope of each trendline at foraging distances (circles with radius 250m, 300m, 350m, 400, 450m, 500m, 750m, 1000m, 1250, 1500m, 1750, or 2000m centred on each study site), *β*_1_ is the partial regression coefficient, and x*_i_* are covariates (habitat variables) at each foraging distance.

We predicted a logarithmic relationship between the number of species representing each functional group and the amount of pollinator habitat available. Our analyses provided robust support for such positive logarithmic relationships between the proportion (amount) of suitable habitat within a landscape and bee species richness across all functional guilds, except for bumblebees (*Bombus* spp.) at foraging distances <500m (Fig. S2d, blue line; Table S5). We did not expect to find a significant logarithmic relationship between the proportional abundance of species in functional guilds and the amount of suitable habitat as this is more likely influenced by site level characteristics (including the availability of nesting and floral resources) at each sampling location – a view supported by previous studies in agricultural (*58*) and urban habitats (*59*). In fact, our results demonstrate that increasing either the availability of nesting or foraging resources at different spatial scales did not support increases in bee abundance for social ground nesters, cavity nesters or bumblebees *Bombus spp.* (Fig. S2g, h, i); but it did for solitary ground nesters and cleptoparasites (Fig. S2f, j).

*2.2 Pollinator habitats within a landscape*

To examine the importance of specific habitat types on pollinator diversity and abundance in various natural and agricultural landscapes, many studies have assessed bee biodiversity metrics (diversity, richness and abundance) at various distances (typically 250, 500, 750, 1000 and 1250 m) from the centre of agricultural fields (e.g., (*48, 52, 60, 61*) to include the flight ranges for most wild bee species estimated from their body size (*62, 63*). For our study, the total extent (amount) of pollinator habitat (defined by Ontario land classes (Tables S3, S4)) was quantified in circles with radii 250, 300, 350, 400, 450, 500, 750, 1000, 1250, 1500, 1750 and 2000 m centred around the 34 survey sites at which bee species were surveyed and collected. Habitat estimates from each of these areas were then pooled into three distinct spatial range categories: <500m, 500-1500m, and >1500m from the survey location.

Pollinator habitat within a landscape at each of the 34 bee survey locations was quantified using the Pollinator Habitat Baseline (PHaB) mapping layer for the province of Ontario. As part of Ontario’s provincial Pollinator Health Action Plan, the Ministry of Natural Resources and Forestry (MNRF) was tasked with the action of ‘assessing land cover in natural habitats, and in agricultural and urban landscapes in southern Ontario to identify and map probable pollinator habitat’. In response, the MNRF created one of the first comprehensive baseline wild pollinator habitat layers (*64*). This layer integrates the Southern Ontario Land Resource Information System 2.0 (SOLRIS) (*65*) and the Annual Crop Inventory (ACI) produced by Agriculture and Agri-Food Canada (AAFC) (*66*), land cover products that provide a ground-truthed comprehensive, standardized landscape level inventory of natural, rural and urban lands at a 15-m resolution (*64*). The creation of the PHaB mapping layer revealed that Southern Ontario has over ~2.7 million hectares of pollinator habitat (approximately 20% of the total area), representing 24 different habitat types (see Ontario Land Classes in Tables S3, S4). However, creating the PHaB mapping layer also revealed that there has been a net loss of nearly 10,000 hectares of pollinator habitat from Ontario over the 10-year period 2002-2012 (*64*). Please refer to Hogg and Jones (*64*) for more information on criteria for selecting quality pollinator habitat in Ontario.

The total area of each pollinator habitat type (Ontario Land Class), defined by PHaB, for each of the twelve nested spatial scales was then quantified using Geographic Information System (GIS) software (*67*). The buffer tool was used to lay down circles with each of the twelve radius distances (range 250-2000 m, described above) centred around each pollinator survey location. The zonal statistics tool was then used to summarise the total extent of each habitat type for each circle, with radii ranging from 250 to 2000 metres. The output habitat area tables were imported into excel, and related back to survey data for analytical purposes. Habitat extents were calculated independently by radius and were not added among the twelve radius distances. So the area closest to each study site was a circle radius 250m, then annulus shaped area with an outer and inner radius of 300 and 250m respectively, etc. As such percentage habitat assessments within each area were only assessed once across all radii from the study site/location.

We tested for correlations among the measured habitat covariates at the maximum foraging distance (2km) using Spearman’s rank correlations (r_s_). Generally, where r_s_ coefficients between two variables exceed 0.7 they are considered to have a strong relationship and should be removed from the analyses (*68*). In our analyses there were correlations between habitat types, but while none of the r_s_ values were higher than 0.7, many were significant (p <0.05: Table S6). The relationship between total pollinator habitat and several other habitat types, were strongly associated (Coniferous Forest: r_s_= 0.49; Deciduous Forest: r_s_= 0.53; Plantation: r_s_= 0.41; Treed Swamp: r_s_ = 0.68, p <0.05; Table S6). A stronger relationship between total pollinator habitat and individual habitat types was expected as total habitat incorporates each individual habitat type. Although the correlation between total pollinator habitat and several habitat types was greater in comparison to other reported relationships (Table S6), we did not remove this parameter (total pollinator habitat), from our analyses. This decision was taken as it was important to test whether the total amount of pollinator habitat was more important for each functional group in comparison to any specific habitat types. A significant relationship between total pollinator habitat and bee functional guilds would suggest that species are more dependent on overall quantity of floral and nesting provisions provided in landscapes rather than provisions provided by specific habitat types.

We analysed which habitats were most important to each of the bee functional guilds using glmm models (candidate model set/functional guild) for species richness and proportional abundance at each radius distance (i.e. 5 functional guilds x 12 radius distances x 2 community parameters (species richness and proportional abundance) = 120 glmms in total; Table S7). Delta AICc (∆AICc), AICc, and AIC weights (*w*) of candidate models were used to select the best fit model within each spatial category (<500m, 500-1500m, >1500m) shown in bold in Table S7. Both functional guild abundance and species richness were log-transformed using ln [a + 1, r + 1] prior to analyses. Although 24 different pollinator habitat types were found in the landscape (Tables S3, S4), we parameterized our global models for each functional guild to test for main effects of the six most influential habitat types on bee parameters. Prioritising the most important (six) habitat types in this way provides end users with a more targeted set of habitat recommendations to meet their specific conservation needs. We did not include interactions between different habitat covariates in our models due to a lack of biological justification for reporting statistical interactions among habitats, that may or may not be within close proximity to one another within a landscape.

Partial regression coefficients (*β_1_*) for habitat covariates (independent variables) used in models of best fit at each spatial category were reported as a way to assess the importance of habitat types for target bee community parameters for each functional guild (Tables S8-10; S8: <500m; S9: 750-1250m; S10: >1500m). We found a number of habitat types were not highlighted as among the six most important for any functional guilds within our spatial categories (Tables S8-10). Abandoned extraction-vegetated, plantations, and treed sand dunes were not found to be important drivers of functional guild species richness or abundance at the most local scales (<500m: Table S8). Abandoned extraction-open and coniferous forest habitats were not found to be important for supporting abundance of any functional guild, whereas built-up pervious habitats were not important parameters for species richness for any guild (Table S8). Our results show that abandoned extractions-open continued to be unimportant for maintaining species richness and abundance of functional guilds at 750-1250m, along with Treed sand dunes (Table S9). Built-up pervious, and total pollinator habitat were not important covariates for any functional guild abundance whereas mixed forests, pastures, and plantations appeared not be important predictors of functional guild species richness in our analyses (Table S9). Tallgrass savannah and treed swamps appeared not to be important predictors for any functional guilds at the largest spatial scales (>1500m: Table S10). Furthermore, at spatial scales >1500m, four different habitats (built-up previous, consistent foraging crop, deciduous forest, and semi-natural) were not selected in any of our best fitting candidate models for functional guild abundance, and two more habitats (marsh and mixed forest) were not selected as supporting species richness for any functional guild (Table S10).

To explore the extent of correlation between preferred habitat types among functional guilds and across spatial scales, we clustered partial regression coefficients of significant habitat types using ascendant hierarchical clustering based on Euclidian distances, then mapped these using a heat map function to visually report results to be easily accessible for all end-users (academics, conservationists, farmers, and policymakers) wanting to interpret the results for conservation and maintenance of habitat for wild bee species (Fig. S3). Colour coding in the heat map indicates the positive (green) and negative (red) impacts of each habitat type for either species richness or proportional abundance for each of the five bee functional guilds. Lighter shades of green indicate a greater importance for the habitat at different spatial distances, where darker shades of red suggest the habitat type is less important for supporting functional guild species richness and abundance. Habitat similarity among functional guilds is characterized by horizontal homogeneous groupings of colours, whereas similar vertical homogenous groupings represent habitat similarities among spatial categories and habitat types (Figs. 4, S3). We used XLSTAT and R 3.0.2 functions to run all statistical analyses (*69, 70*).

**Supplementary tables and figures**

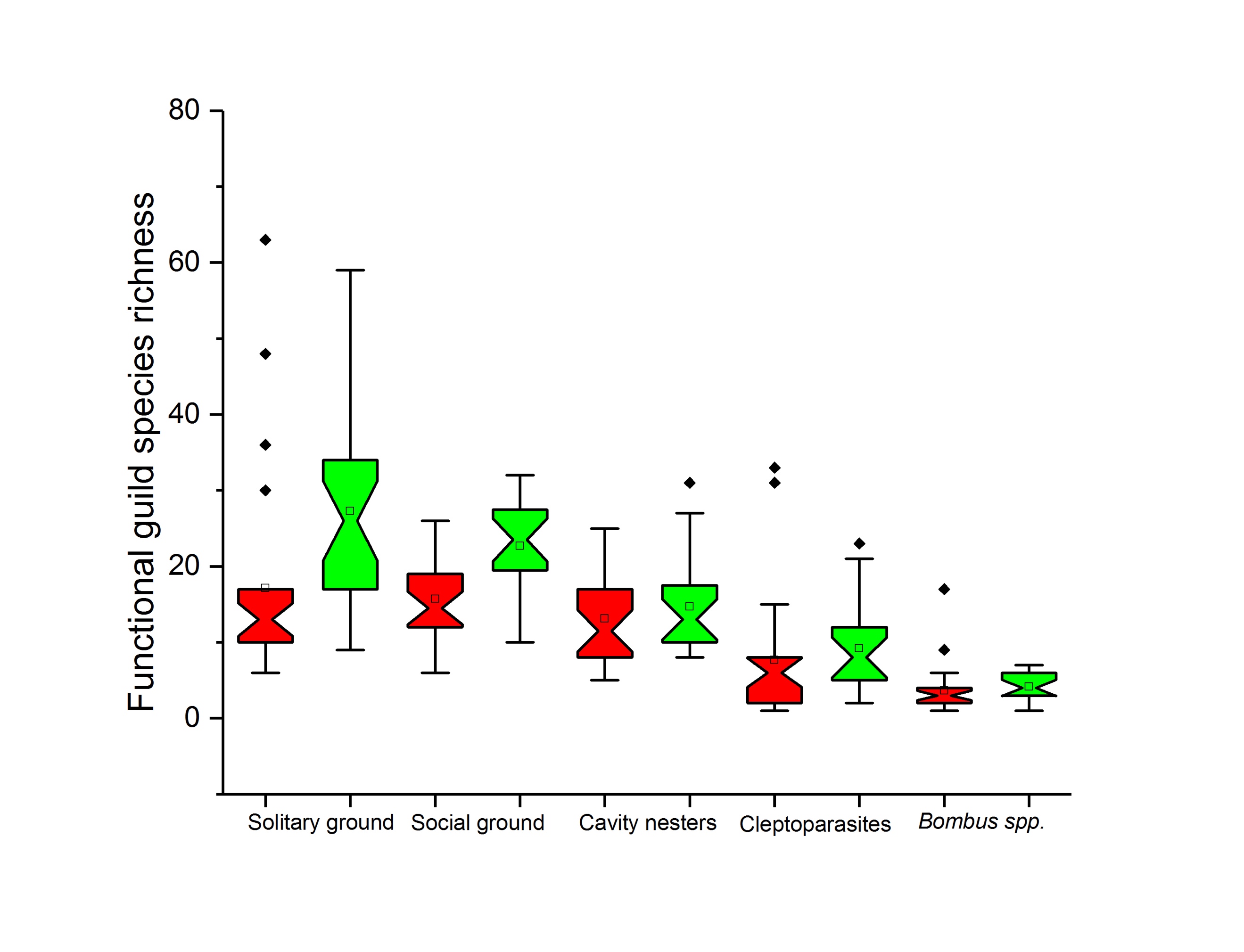
 (a)

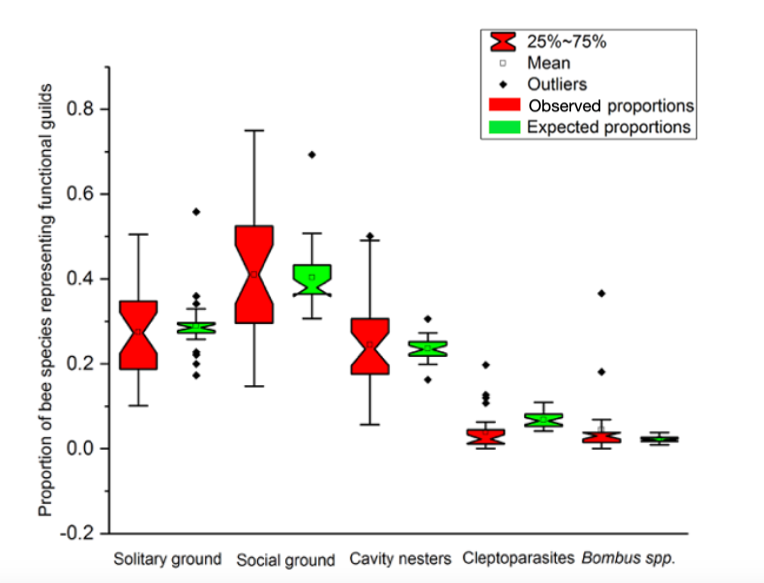
 (b)

Fig S1. Notched-box plots depicting differences between observed and expected (a) species richness and (b) proportion of bee species representing each functional guild: solitary ground nesters; social ground nesters; cavity nesters; cleptoparasites; and *Bombus* spp. Alignment of notches denotes no significant between observed and expected species richness and proportional abundance for each functional guild.

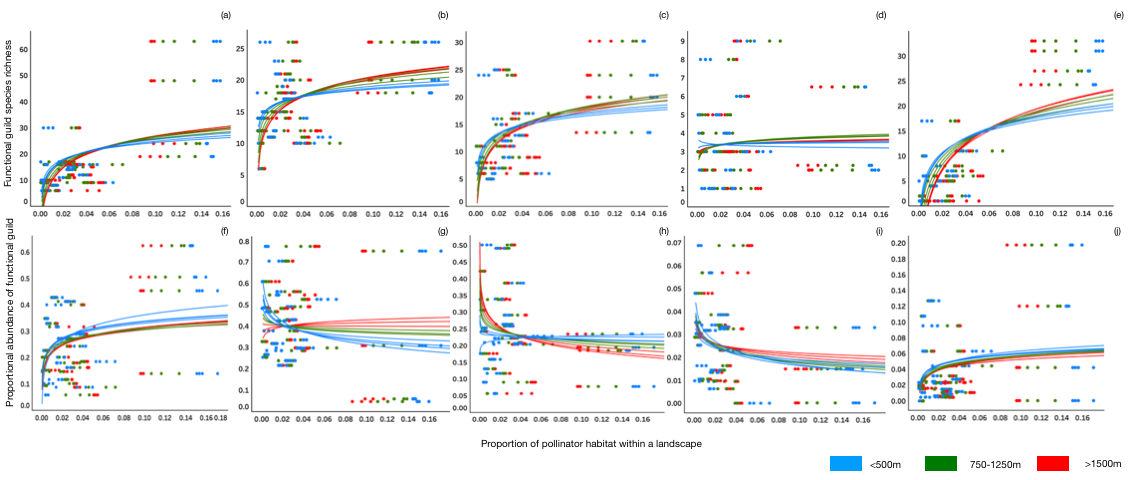

Fig S2. Relationship between the proportion of pollinator habitat within a landscape in Southern Ontario and community parameters, species richness (top row) and proportional abundance (bottom row) of solitary ground nesting bee species (a,f); social ground nesters (b,g); cavity nesters (c,h); *Bombus* spp. (d,i); cleptoparasites (e, j). Logarithmic trendlines in blue are for foraging distances <500m, green trendlines represent foraging distances between 750-1250m, and red trendlines are distances >1500m. Regression coefficients and p values associated with each spatial scale trendline and functional guild community parameter can be found in Table S5.

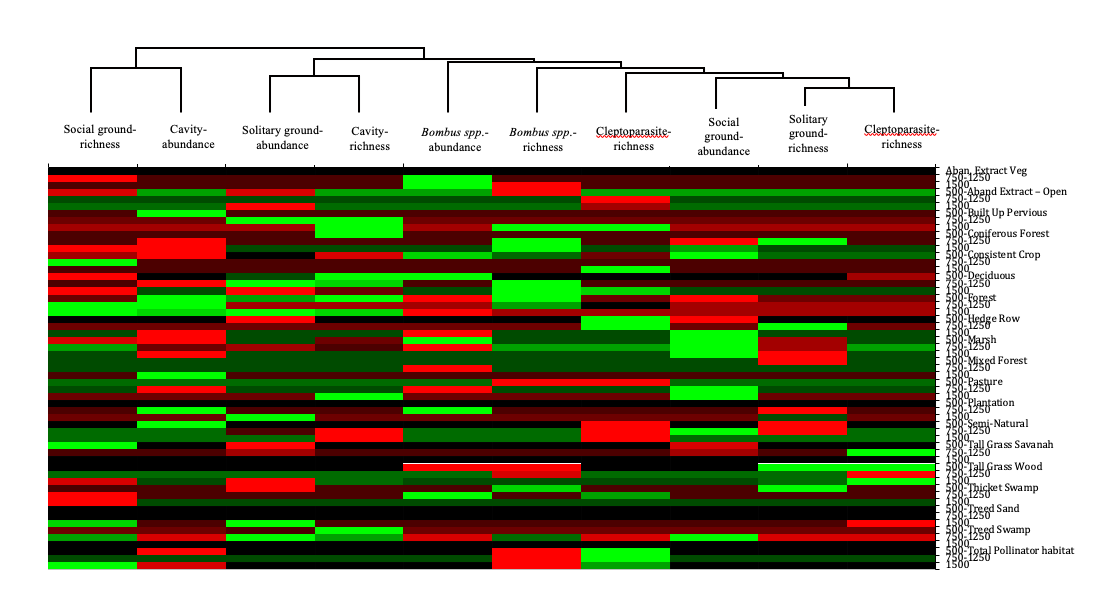

Fig S3. Heat map of habitat model parameters and their importance for maintaining expected bee biodiversity metrics: species richness and proportional abundance (Fig. 2) of functional guilds: solitary ground nesters, social ground nesters, cavity nesters, *Bombus* spp., and cleptoparasitic species at foraging distance categories (<500m, between 750-1500m, and >1500m). Lighter shades of green indicate a greater importance for the habitat at different spatial distances, where darker shades of red suggest a less desirable habitat for supporting functional guild species richness and abundance. Habitat similarity is characterized by groupings of a colour: among function guilds (horizontal rows), and across spatial distances and habitat types (vertical columns). Forested habitats represented 50m of habitat edges. Black cells imply the habitat has a neutral impact on bee species richness and/or abundance in the landscape.

Table S1. Distribution of the 361 wild bee species (from 66,271 individual records) among 39 genera and comparisons of database generic richness with the Ontario bee species list generated from published and unpublished studies (*36, 71-77*).

| Family | Genus | Number of species | | Percentage of Ontario species in database |
| --- | --- | --- | --- | --- |
|  |  | Database | Ontario |  |
| Andrenidae | *Andrena* spp. | 64 | 75 | 85% |
|  | *Calliopsis* spp. | 1 | 1 | 100% |
|  | *Perdita* spp. | 4 | 6 | 67% |
|  | *Pseudopanurgus* spp. | 2 | 2 | 100% |
| Apidae | *Anthophora* spp. | 2 | 2 | 100% |
|  | *Bombus* spp. | 19 | 25 | 76% |
|  | *Ceratina* spp. | 4 | 4 | 100% |
|  | *Epeolus* spp. | 8 | 12 | 67% |
|  | *Eucera spp.* | 1 | 1 | 100% |
|  | *Holcopasites* spp. | 1 | 1 | 100% |
|  | *Melissodes* spp. | 11 | 11 | 100% |
|  | *Nomada* spp. | 34 | 34 | 100% |
|  | *Ptilothrix* spp. | 0 | 1 | - |
|  | *Svastra* spp. | 0 | 1 | - |
|  | *Triepeolus* spp. | 2 | 9 | 22% |
|  | *Xylocopa* spp. | 1 | 1 | 100% |
| Colletidae | *Colletes* spp. | 4 | 17 | 24% |
|  | *Hylaeus* spp. | 12 | 14 | 86% |
| Halictidae | *Agapostemon* spp. | 4 | 4 | 100% |
|  | *Augochlora* spp. | 1 | 1 | 100% |
|  | *Augochlorella* spp. | 1 | 1 | 100% |
|  | *Augochloropsis* spp. | 1 | 1 | 100% |
|  | *Dieunomia* spp. | 0 | 1 | - |
|  | *Duforea* spp. | 0 | 5 | - |
|  | *Halictus* spp. | 4 | 4 | 100% |
|  | *Lasioglossum* spp. | 66 | 74 | 89% |
|  | *Sphecodes* spp. | 22 | 23 | 96% |
| Megachilidae | *Anthidellum* spp. | 1 | 1 | 100% |
|  | *Anthidium* spp. | 1 | 3 | 33% |
|  | *Chelostoma* spp. | 1 | 3 | 33% |
|  | *Coelioxys* spp. | 3 | 10 | 30% |
|  | *Chelostoma* spp. | 1 | 3 | 33% |
|  | *Dianthidium* spp. | 0 | 1 | - |
|  | *Heriades* spp. | 3 | 3 | 100% |
|  | *Hoplitis* spp. | 5 | 7 | 71% |
|  | *Megachile* spp. | 18 | 20 | 90% |
|  | *Osmia* spp. | 22 | 24 | 92% |
|  | *Stelis* spp. | 4 | 8 | 50% |
| Melittidae | *Macropis* spp. | 1 | 1 | 100% |
|  | **Total** | **361** | **421** | **86%** |

Table S2. Mood test of observed and null (expected) bee community parameters: a) species richness and b) proportional abundance. Data presented are the expected (Exp.) and observed (Obs.) values for each parameter, the percentage difference between observed and expected values, the U statistic (critical value = 3.84) and p values for these comparisons. Significant comparisons are shown in bold.

| Functional guild | a) Species richness | | | | | b) Proportional abundance | | | | |
| --- | --- | --- | --- | --- | --- | --- | --- | --- | --- | --- |
|  | Exp. | Obs. | % diff | U | p | Exp. | Obs. | % diff | U | p |
| Solitary ground | **17.15** | **27.27** | **59.01** | **17.43** | **<0.0001** | 0.29 | 0.27 | -6.90 | 0.016 | 0.8984 |
| Social ground | **15.13** | **22.70** | **50.03** | **15.01** | **<0.0001** | 0.42 | 0.41 | -2.38 | 0.019 | 0.8910 |
| Cavity nesters | 13.11 | 14.70 | 12.13 | 1.143 | 0.2850 | 0.22 | 0.25 | -13.64 | 0.018 | 0.8808 |
| *Bombus* spp. | 3.65 | 4.16 | 13.97 | 1.523 | 0.2127 | 0.04 | 0.04 | 0.00 | 0.026 | 0.8991 |
| Cleptoparasites | **7.68** | **9.17** | **19.40** | **13.24** | **0.0002** | 0.03 | 0.04 | +33.33 | 0.027 | 0.8913 |

Table S3. Ontario land classes found within 2 km of each of 34 bee sampling locations in Southern Ontario. Total amounts of each habitat found in sampling location and total amount in Southern Ontario from Hogg and Jones (*64*).

| **Ontario Land Classes** | **Pollinator provisions from habitat** | **Total amount in Southern Ontario (ha)** | **Habitat percentage of out of total** | **Total habitat in surveys (at 2km) (ha)** | **Habitat in surveys percentage (%)** |
| --- | --- | --- | --- | --- | --- |
| Abandoned Extraction Open | Nesting | 22940 | 0.83 | 1.88 | 1.07 |
| Abandoned Extraction Vegetated | Nesting | 4374 | 0.16 | 0.78 | 0.45 |
| Built Up Area Pervious (e.g. veg. parks) | Forage & nesting | 91180 | 3.28 | 9.66 | 5.52 |
| Coniferous Forest | Nesting | 234475 | 8.44 | 8.69 | 4.96 |
| Consistent Forage crop (AAFC land data) | Forage | 52509 | 1.89 | 2.25 | 1.29 |
| Deciduous Forest | Nesting | 581021 | 20.90 | 39.01 | 22.29 |
| Forest | Nesting | 45672 | 1.64 | 2.41 | 1.38 |
| Hedge Row | Forage & nesting | 56741 | 2.04 | 7.28 | 4.16 |
| Marsh | Forage & nesting | 141153 | 5.08 | 4.83 | 2.76 |
| Mixed Forest | Nesting | 269536 | 9.70 | 21.2 | 12.12 |
| Open Alvar | Forage & nesting | 2039 | 0.07 | - | - |
| Open Cliff and Talus | Forage | 20 | 0.00 | - | - |
| Open Sand Dune | Nesting | 698 | 0.03 | - | - |
| Open Tallgrass Prairie | Forage & nesting | 295 | 0.01 | - | - |
| Pasture | Forage & Nesting | 150000 | 5.40 | 0.23 | 0.01 |
| Plantation | Forage & nesting | 87154 | 3.14 | 6.83 | 3.9 |
| Semi-Natural | Forage & nesting | 72238 | 2.60 | 5.29 | 3.02 |
| Shrub Alvar | Forage & nesting | 699 | 0.03 | - | - |
| Tallgrass Savannah | Forage & nesting | 694 | 0.02 | 0.04 | 0.02 |
| Tallgrass Woodland | Forage & nesting | 1208 | 0.04 | 12.23 | 6.99 |
| Thicket Swamp | Nesting | 118904 | 4.28 | 2.68 | 1.53 |
| Treed Alvar | Forage & nesting | 545 | 0.02 | - | - |
| Treed Cliff and Talus | Forage | 125 | 0.00 | - | - |
| Treed Sand Dune | Forage & nesting | 503 | 0.02 | 0.75 | 0.43 |
| Treed Swamp | Nesting | 844695 | 30.39 | 49.2 | 28.11 |
| **Totals** |  | 2,779,418 (ha) | 100% | 175.24 (ha) | 100% |

Table S4. Description of the Ontario habitat types found within 2 km of each bee sampling location as determined by Lee (*78*) and Hogg and Jones (*64*).

| **Ontario Land Classes/ Habitat Type** | **Plant Community** |
| --- | --- |
| Abandoned Extraction Open | Inactive pit/quarry |
| Abandoned Extraction Vegetated | Inactive pit/quarry with tree cover ≤ 25%; shrub cover ≤ 25% |
| Built Up Area Pervious (e.g. veg. parks) | Vegetated Park area |
| Coniferous Forest | Largely continuous forest canopy composed primarily of coniferous species; includes swamps. |
| Consistent Forage crop (AAFC land data) | Open Agriculture- annual row crops Tobacco, ginseng, oilseeds, borage, camelina, canola/rapeseed, flaxseed, mustard, safflower, sunflower, soybeans, pulses, peas, beans, lentils, vegetables, tomatoes, potatoes, sugar beets, other vegetables, fruits, berries, cranberry, orchards, other fruits, vineyards, hops, nursery, buckwheat, vetch, fallow |
| Deciduous Forest | Deciduous tree species > 75% of canopy cover |
| Forest | Tree cover > 60% |
| Hedge Row | Shrub agriculture land |
| Marsh | A wetland with a mineral or peat substrate inundated by nutrient-rich water and characterized by emergent graminoid vegetation |
| Mixed Forest | forest canopy composed of both deciduous and coniferous species, - conifer tree species > 25% and deciduous tree species > 25% of canopy cover |
| Open Tallgrass Prairie | Natural areas typically have unique floras (e.g. Tallgrass Savannah), areas with a cultural legacy, typically dominated by more invasive herbaceous, shrub, and tree species; tree cover typically scattered or clumped |
| Pasture | Field dominated with herbaceous vegetation and grasses with an understory of similar material in a state of decay |
| Plantation | Treed agriculture land |
| Semi-Natural | Area containing natural plant communities but with obvious human alteration in form of maintenance |
| Tallgrass Savannah | 25% < tree cover < 35%; semi-open treed communities; natural areas typically have unique floras (e.g. Tallgrass Savannah), areas with a cultural legacy, typically dominated by more invasive herbaceous, shrub, and tree species; tree cover typically scattered or clumped |
| Tallgrass Woodland | 35% < tree cover < 60%; semi-closed treed communities; natural areas typically have unique floras (e.g. Tallgrass Woodland), areas with a cultural legacy, typically dominated by more invasive herbaceous, shrub, and tree species; tree cover more closed and shaded |
| Thicket Swamp | Tree or shrub cover > 25% - dominated by hydrophytic shrub and tree species |
| Treed Sand Dune | Tree cover > 25% tree cover varies from scattered and clumped to more continuous cover |
| Treed Swamp | A mineral-rich wetland characterized by a dense cover of deciduous or coniferous trees or shrubs - tree cover > 25%; trees > 5 m in height - deciduous tree species > 75% of canopy cover - common species include Fowl Manna Grass, Spotted Touch -me -not, Bugleweed, Skunk Cabbage, Marsh Marigold, Bedstraws and Stinging Nettles - typically fern and sedge rich |

Table S5. Regression results for relationships between pollinator habitat within a landscape and bee community, parameters species richness and proportional abundance, for each of the five functional guilds: solitary ground nesters, social ground nesters, cavity nesters, *Bombus* spp. and cleptoparasites. F is the foraging distance (m) from each site within the landscape. Significant results are shown in bold (alpha = 0.05), p values <0.005 represented by *** due to space constraints in table.

| F/ (m) | Solitary ground | | | | | | | Social ground | | | | | | | | Cavity | | | | | | | | | | *Bombus* spp. | | | | Cleptoparasite | | | | |
| --- | --- | --- | --- | --- | --- | --- | --- | --- | --- | --- | --- | --- | --- | --- | --- | --- | --- | --- | --- | --- | --- | --- | --- | --- | --- | --- | --- | --- | --- | --- | --- | --- | --- | --- |
|  | richness | | | pr.abundance | | | | richness | | | pr.abundance | | | | richness | | | | pr abundance | | | | | | richness | | | pr.abundance | | richness | | | pr.abundance | |
|  | R^2^ | | p | R^2^ | | *p* | | R^2^ | | p | | R^2^ | | *p* | R^2^ | | | p | | R^2^ | | | *p* | | R^2^ | | p | R^2^ | *p* | R^2^ | p | | R^2^ | *p* |
| 250 | **0.26** | **0.02** | | | 0.13 | | 0.12 | 0.14 | 0.11 | | | | 0.005 | 0.76 | 0.05 | | 0.34 | | | | 0.11 | 0.15 | | 0.01 | | | 0.67 | 0.06 | 0.51 | **0.37** | | **0.01** | 0.08 | 0.11 |
| 300 | **0.22** | **0.04** | | | 0.08 | | 0.39 | 0.14 | 0.17 | | | | <0.001 | 0.98 | 0.09 | | 0.20 | | | | 0.13 | 0.13 | | <0.001 | | | 0.85 | 0.002 | 0.92 | **0.39** | | ******* | **0.04** | 0.24 |
| 350 | **0.08** | **0.19** | | | 0.02 | | 0.54 | 0.07 | 0.25 | | | | 0.022 | 0.52 | **0.25** | | **0.02** | | | | <0.001 | 0.89 | | <0.001 | | | 0.97 | 0.002 | 0.68 | **0.11** | | **0.14** | 0.001 | 0.97 |
| 400 | **0.30** | **0.01** | | | 0.02 | | 0.45 | **0.29** | **0.01** | | | | 0.05 | 0.30 | **0.39** | | ******* | | | | 0.002 | 0.87 | | 0.10 | | | 0.16 | 0.01 | 0.53 | **0.42** | | ******* | <0.001 | 0.60 |
| 450 | **0.29** | **0.01** | | | 0.03 | | 0.39 | **0.25** | **0.01** | | | | 0.062 | 0.22 | **0.44** | | ******* | | | | 0.003 | 0.81 | | 0.03 | | | 0.40 | 0.01 | 0.51 | **0.48** | | ******* | 0.017 | 0.30 |
| 500 | **0.29** | **0.01** | | | 0.03 | | 0.44 | **0.24** | **0.01** | | | | 0.0004 | 0.92 | **0.36** | | ******* | | | | 0.04 | 0.31 | | 0.009 | | | 0.65 | 0.03 | 0.33 | **0.44** | | ******* | 0.012 | 0.28 |
| 750 | **0.23** | **0.01** | | | 0.07 | | 0.14 | **0.22** | **0.02** | | | | 0.006 | 0.78 | **0.36** | | **0.04** | | | | **0.23** | **0.01** | | <0.001 | | | 0.95 | 0.002 | 0.71 | **0.38** | | ******* | 0.05 | 0.09 |
| 1000 | **0.29** | **0.01** | | | 0.07 | | 0.12 | **0.32** | ******* | | | | 0.04 | 0.40 | **0.17** | | **0.06** | | | | **0.18** | **0.02** | | 0.025 | | | 0.44 | 0.03 | 0.46 | **0.33** | | ******* | 0.09 | 0.20 |
| 1250 | **0.24** | **0.01** | | | 0.05 | | 0.23 | **0.27** | **0.01** | | | | 0.05 | 0.31 | **0.14** | | **0.07** | | | | 0.10 | 0.09 | | 0.025 | | | 0.44 | 0.012 | 0.69 | **0.33** | | ******* | <0.001 | 0.32 |
| 1500 | **0.22** | **0.01** | | | 0.02 | | 0.41 | **0.18** | **0.03** | | | | 0.09 | 0.15 | 0.13 | | 0.18 | | | | **0.25** | **0.01** | | 0.013 | | | 0.58 | 0.003 | 0.96 | **0.30** | | **0.01** | 0.003 | 0.41 |
| 1750 | **0.14** | **0.05** | | | 0.001 | | 0.73 | 0.10 | 0.11 | | | | 0.14 | 0.07 | 0.07 | | 0.31 | | | | **0.31** | ******* | | 0.008 | | | 0.67 | <0.001 | 0.92 | **0.19** | | **0.03** | 0.004 | 0.73 |
| 2000 | **0.27** | **0.01** | | | 0.05 | | 0.24 | **0.28** | **0.01** | | | | 0.01 | 0.62 | 0.04 | | 0.01 | | | | 0.16 | 0.08 | | 0.003 | | | 0.78 | 0.003 | 0.62 | **0.42** | | ******* | 0.004 | 0.29 |

| Ontario Land Classes/ Habitat Type | Ab.Ex. Open | Ab.Ex. Veget. | Bu. Up Area | Conif. Forest | Cons F Crop | Decid. Forest | Forest | Hedge Row | Marsh | Mixed Forest | Open Beach | Pasture | Plant. | Semi-Natural | Tallgr Savann | Tallgr Wood | Thick.Swamp | Tr. San Dune | Tr. Swamp. | Tot Pol Habitat |
| --- | --- | --- | --- | --- | --- | --- | --- | --- | --- | --- | --- | --- | --- | --- | --- | --- | --- | --- | --- | --- |
| Abandoned Extraction Open | **1.00** |  |  |  |  |  |  |  |  |  |  |  |  |  |  |  |  |  |  |  |
| Abandoned Extraction Vegetated | 0.01 | **1.00** |  |  |  |  |  |  |  |  |  |  |  |  |  |  |  |  |  |  |
| Built Up Area Pervious | 0.12 | 0.17 | **1.00** |  |  |  |  |  |  |  |  |  |  |  |  |  |  |  |  |  |
| Coniferous Forest | 0.01 | 0.01 | **0.21** | **1.00** |  |  |  |  |  |  |  |  |  |  |  |  |  |  |  |  |
| Consistent Foraging Crop | 0.01 | 0.01 | 0.10 | 0.01 | **1.00** |  |  |  |  |  |  |  |  |  |  |  |  |  |  |  |
| Deciduous Forest | 0.03 | 0.05 | **0.32** | **0.37** | 0.05 | **1.00** |  |  |  |  |  |  |  |  |  |  |  |  |  |  |
| Forest | 0.01 | 0.01 | **0.20** | **0.31** | 0.04 | **0.40** | **1.00** |  |  |  |  |  |  |  |  |  |  |  |  |  |
| Hedge Row | **0.16** | 0.00 | **0.36** | **0.26** | 0.04 | **0.31** | 0.14 | **1.00** |  |  |  |  |  |  |  |  |  |  |  |  |
| Marsh | 0.02 | 0.05 | **0.29** | **0.38** | 0.02 | **0.46** | **0.18** | **0.24** | **1.00** |  |  |  |  |  |  |  |  |  |  |  |
| Mixed Forest | 0.01 | 0.01 | 0.05 | **0.20** | 0.06 | 0.06 | **0.35** | 0.00 | **0.17** | **1.00** |  |  |  |  |  |  |  |  |  |  |
| Open Beach Bar | 0.05 | 0.01 | 0.04 | 0.03 | 0.00 | 0.10 | 0.02 | 0.10 | 0.00 | 0.00 | **1.00** |  |  |  |  |  |  |  |  |  |
| Pasture | 0.07 | 0.04 | 0.06 | 0.04 | **0.22** | 0.00 | 0.01 | 0.04 | 0.01 | 0.01 | **0.19** | **1.00** |  |  |  |  |  |  |  |  |
| Plantation | 0.03 | 0.11 | **0.24** | **0.22** | 0.03 | **0.31** | 0.06 | 0.06 | **0.16** | 0.15 | 0.11 | 0.00 | **1.00** |  |  |  |  |  |  |  |
| Semi-Natural | 0.02 | 0.00 | 0.01 | **0.04** | **0.18** | 0.01 | 0.01 | **0.15** | 0.01 | 0.02 | **0.16** | **0.24** | 0.01 | **1.00** |  |  |  |  |  |  |
| Tallgrass Savanah | 0.14 | 0.01 | 0.01 | 0.00 | 0.02 | 0.11 | 0.05 | 0.10 | 0.04 | 0.11 | **0.52** | 0.09 | 0.01 | 0.08 | **1.00** |  |  |  |  |  |
| Tallgrass Woodland | 0.05 | 0.01 | 0.04 | 0.03 | 0.00 | 0.10 | 0.02 | 0.10 | 0.00 | 0.00 | 0.48 | **0.19** | 0.11 | **0.16** | 0.52 | **1.00** |  |  |  |  |
| Thicket Swamp | 0.28 | 0.00 | **0.39** | **0.44** | 0.00 | **0.51** | **0.21** | **0.49** | **0.47** | 0.02 | 0.01 | 0.04 | 0.14 | 0.00 | 0.02 | 0.01 | **1.00** |  |  |  |
| Treed Sand Dune | 0.14 | 0.01 | 0.01 | 0.00 | 0.02 | 0.11 | 0.05 | 0.10 | 0.04 | 0.11 | **0.52** | 0.09 | 0.01 | 0.08 | **0.56** | 0.52 | 0.02 | **1.00** |  |  |
| Treed Swamp | 0.00 | 0.00 | 0.01 | **0.26** | 0.03 | **0.16** | **0.26** | 0.00 | 0.12 | **0.25** | 0.00 | **0.34** | **0.17** | 0.02 | 0.01 | 0.00 | 0.13 | **0.01** | **1.00** |  |
| Total Pollinator Habitat | 0.04 | 0.00 | **0.15** | **0.49** | 0.00 | **0.53** | **0.35** | 0.10 | **0.33** | **0.18** | 0.15 | **0.15** | **0.41** | 0.00 | 0.08 | 0.15 | **0.42** | **0.08** | **0.68** | **1.00** |

Table S6. Matrix of Spearman’s Rank correlation coefficients of relationship between habitat types. Coefficients of determination greater than 0.7 are considered strong correlations at a foraging distance of 2000m. Values in bold are different from 0 with a significance level p<0.05. Values shown as 0.00 indicate coefficients <0.005. Grey shaded cells are self-correlated habitat types (coefficient = 1.00).

Table S7. Candidate models testing relationships between bee functional guild (solitary ground nesters, social ground nesters, cavity nesters, *Bombus* spp. and cleptoparasites) measures (species richness and proportional abundance) and pollinator habitats (model variables) in landscape and model selection statistics. F is the foraging distance (m) from each site within the landscape; K is the number of parameters included in each candidate model; model parameters are habitat types, and AIC is Akaike’s Information Criterion, and *w* is AIC weight. Candidate models in bold are considered to be models of best fit in each spatial category (<500m; 750-1250m; >1500m).

| **Solitary Ground Species Richness** | | | |  |  |
| --- | --- | --- | --- | --- | --- |
| **F** | **K** | **Model Variables (Ontario Land Classes/ Habitat Types)** | **AIC** | ΔAIC | *w* |
| 250 | 3 | Treed Swamp / Consistent Forage Crop / Tallgrass Woodland | -48.9 | 13.5 | 0.00 |
| 300 | 3 | Treed Swamp / Consistent Forage Crop / Tallgrass Woodland | -49.3 | 13.1 | 0.00 |
| 350 | 3 | Treed Swamp / Consistent Forage Crop / Tallgrass Woodland | -49.3 | 13.2 | 0.00 |
| 400 | 3 | Treed Swamp / Consistent Forage Crop / Tallgrass Woodland | -54.2 | 8.2 | 0.00 |
| 450 | 6 | Treed Swamp / Forest / Coniferous Forest / Built Up Area Pervious / Consistent Forage Crop / Tallgrass Savannah | -62.0 | 0.4 | 0.01 |
| **500** | **6** | **Treed Swamp / Forest / Deciduous Forest / Thicket Swamp / Consistent Forage Crop / Tallgrass Savannah** | **-62.5** | **0.0** | **0.02** |
| 750 | 6 | Treed Swamp / Forest / Deciduous Forest / Marsh / Abandoned Extraction Open / Tallgrass Savannah | -70.5 | 1.9 | 0.09 |
| **1000** | **6** | **Treed Swamp / Deciduous Forest / Marsh / Semi-Natural/ Built Up Area Pervious / Tallgrass Savannah** | **-72.4** | **0.0** | **0.25** |
| 1250 | 6 | Treed Swamp / Coniferous Forest / Mixed Forest / Deciduous Forest / Marsh / Built Up Area Pervious | -70.5 | 1.9 | 0.09 |
| 1500 | 6 | Forest / Mixed Forest / Plantation / Abandoned Extraction Open / Tallgrass Woodland / Tallgrass Savannah | -69.6 | 4.1 | 0.02 |
| **1750** | **6** | **Forest / Deciduous Forest / Plantation / Abandoned Extraction Open / Tallgrass Woodland / Treed Sand Dune** | **-73.7** | **0.0** | **0.18** |
| 2000 | 6 | Forest / Coniferous Forest / Mixed Forest / Plantation / Tallgrass Woodland / Treed Sand Dune | -72.3 | 1.4 | 0.09 |
| **Social Ground Species Richness** | | | |  |  |
| 250 | 6 | Deciduous Forest / Marsh / Hedge Row / Abandoned Extraction Open / Consistent Forage Crop / Tallgrass Savannah | -58.8 | 8.4 | 0.01 |
| 300 | 6 | Deciduous Forest / Marsh / Hedge Row / Abandoned Extraction Open / Consistent Forage Crop / Tallgrass Savannah | -62.3 | 4.9 | 0.06 |
| **350** | **6** | **Deciduous Forest / Marsh / Hedge Row / Abandoned Extraction Open / Consistent Forage Crop / Tallgrass Savannah** | **-67.2** | **0.0** | **0.64** |
| 400 | 2 | Built Up Area Pervious / Total Pollinator Habitat | -57.1 | 10.2 | 0.00 |
| 450 | 6 | Pasture / Forest / Plantation / Built Up Area Pervious / Consistent Forage Crop / Total Pollinator Habitat | -63.9 | 3.3 | 0.12 |
| 500 | 6 | Forest / Marsh / Built Up Area Pervious / Abandoned Extraction Open / Consistent Forage Crop / Total Pollinator Habitat | -64.6 | 2.6 | 0.17 |
| 750 | 5 | Forest / Marsh / Built Up Area Pervious / Abandoned Extraction Open / Total Pollinator Habitat | -75.9 | 4.2 | 0.10 |
| 1000 | 6 | Forest / Coniferous Forest / Abandoned Extraction Open / Abandoned Extraction Vegetated / Tallgrass Savannah / Total Pollinator Habitat | -73.8 | 6.3 | 0.04 |
| **1250** | **6** | **Treed Swamp / Forest / Thicket Swamp / Abandoned Extraction Open / Consistent Forage Crop / Abandoned Extraction Vegetated** | **-80.1** | **0.0** | **0.86** |
| 1500 | 6 | Forest / Coniferous Forest / Deciduous Forest / Marsh / Abandoned Extraction Open / Total Pollinator Habitat | -74.2 | 5.0 | 0.06 |
| 1750 | 6 | Treed Swamp / Forest / Coniferous Forest / Deciduous Forest / Thicket Swamp / Total Pollinator Habitat | -77.4 | 1.9 | 0.26 |
| **2000** | **5** | **Forest / Coniferous Forest / Deciduous Forest / Thicket Swamp / Total Pollinator Habitat** | **-79.2** | **0.0** | **0.68** |
| **Cavity Nesters Species Richness** | | | |  |  |
| 250 | 5 | Treed Swamp / Pasture / Built Up Area Pervious / Abandoned Extraction Open / Tallgrass Savannah | -44.4 | 24.0 | 0.00 |
| 300 | 5 | Forest / Deciduous Forest / Abandoned Extraction Open / Consistent Forage Crop / Total Pollinator Habitat | -43.4 | 25.0 | 0.00 |
| 350 | 5 | Treed Swamp / Built Up Area Pervious / Abandoned Extraction Open / Consistent Forage Crop / Tallgrass Savannah | -45.0 | 23.4 | 0.00 |
| 400 | 4 | Deciduous Forest / Abandoned Extraction Open / Consistent Forage Crop / Total Pollinator Habitat | -47.3 | 21.1 | 0.00 |
| 450 | 6 | Treed Swamp / Pasture / Forest / Coniferous Forest / Built Up Area Pervious / Consistent Forage Crop | -58.4 | 10.0 | 0.01 |
| **500** | **6** | **Treed Swamp / Forest / Deciduous Forest / Marsh / Abandoned Extraction Open / Consistent Forage Crop** | **-68.4** | **0.0** | **0.99** |
| 750 | 6 | Forest / Mixed Forest / Deciduous Forest / Marsh / Plantation / Consistent Forage Crop | -71.1 | 1.5 | 0.21 |
| 1000 | 6 | Treed Swamp / Mixed Forest / Deciduous Forest / Plantation / Built Up Area Pervious / Abandoned Extraction Vegetated | -72.2 | 0.4 | 0.35 |
| **1250** | **5** | **Treed Swamp / Deciduous Forest / Marsh / Semi-Natural/ Built Up Area Pervious** | **-72.6** | **0.0** | **0.44** |
| 1500 | 6 | Mixed Forest / Deciduous Forest / Plantation / Semi-Natural/ Built Up Area Pervious / Abandoned Extraction Vegetated | -69.2 | 4.8 | 0.05 |
| 1750 | 6 | Pasture / Forest / Semi-Natural/ Built Up Area Pervious / Abandoned Extraction Vegetated / Tallgrass Woodland | -73.6 | 0.4 | 0.43 |
| **2000** | **6** | **Pasture / Forest / Deciduous Forest / Semi-Natural/ Built Up Area Pervious / Tallgrass Woodland** | **-73.9** | **0.0** | **0.52** |
| ***Bombus* Species Richness** | | | |  |  |
| 250 | 3 | Deciduous Forest / Marsh / Built Up Area Pervious | -36.4 | 27.3 | 0.00 |
| 300 | 2 | Forest / Deciduous Forest | -37.0 | 26.7 | 0.00 |
| 350 | 5 | Pasture / Coniferous Forest / Marsh / Hedge Row / Total Pollinator Habitat | -38.3 | 25.4 | 0.00 |
| 400 | 6 | Coniferous Forest / Deciduous Forest / Marsh / Hedge Row / Tallgrass Woodland / Total Pollinator Habitat | -43.0 | 20.7 | 0.00 |
| 450 | 4 | Pasture / Forest / Coniferous Forest / Tallgrass Woodland | -49.0 | 14.7 | 0.00 |
| **500** | **6** | **Pasture / Forest / Coniferous Forest / Thicket Swamp / Tallgrass Woodland / Total Pollinator Habitat** | **-63.7** | **0.0** | **1.00** |
| **750** | **6** | **Treed Swamp / Forest / Coniferous Forest / Deciduous Forest / Tallgrass Woodland / Total Pollinator Habitat** | **-64.2** | **0.0** | **0.85** |
| 1000 | 4 | Deciduous Forest / Built Up Area Pervious / Consistent Forage Crop / Total Pollinator Habitat | -59.2 | 5.0 | 0.07 |
| 1250 | 6 | Pasture / Deciduous Forest / Marsh / Hedge Row / Abandoned Extraction Vegetated / Total Pollinator Habitat | -59.6 | 4.6 | 0.09 |
| 1500 | 5 | Coniferous Forest / Deciduous Forest / Built Up Area Pervious / Tallgrass Savannah / Total Pollinator Habitat | -65.1 | 0.4 | 0.25 |
| **1750** | **6** | **Coniferous Forest / Deciduous Forest / Built Up Area Pervious / Abandoned Extraction Vegetated / Treed Sand Dune / Total Pollinator Habitat** | **-66.2** | **0.0** | **0.31** |
| **2000** | **6** | **Coniferous Forest / Deciduous Forest / Built Up Area Pervious / Abandoned Extraction Vegetated / Treed Sand Dune / Total Pollinator Habitat** | **-65.5** | **0.0** | **0.31** |
| **Cleptoparasite Species Richness** | | | |  |  |
| 250 | 6 | Treed Swamp / Pasture / Deciduous Forest / Semi-Natural/ Abandoned Extraction Open / Tallgrass Woodland | -64.4 | 2.8 | 0.14 |
| 300 | 5 | Pasture / Semi-Natural/ Abandoned Extraction Open / Consistent Forage Crop / Total Pollinator Habitat | -63.1 | 4.1 | 0.07 |
| **350** | **5** | **Pasture / Semi-Natural/ Abandoned Extraction Open / Consistent Forage Crop / Total Pollinator Habitat** | **-67.2** | **0.0** | **0.58** |
| 400 | 6 | Treed Swamp / Forest / Coniferous Forest / Semi-Natural/ Abandoned Extraction Open / Consistent Forage Crop | -58.3 | 8.8 | 0.01 |
| 450 | 6 | Forest / Semi-Natural/ Built Up Area Pervious / Abandoned Extraction Open / Consistent Forage Crop / Total Pollinator Habitat | -65.1 | 2.1 | 0.20 |
| 500 | 6 | Treed Swamp / Forest / Deciduous Forest / Thicket Swamp / Plantation / Abandoned Extraction-Open | -57.4 | 9.7 | 0.00 |
| **750** | **6** | **Forest / Thicket Swamp / Hedge Row / Semi-Natural/ Abandoned Extraction-Open / Total Pollinator Habitat** | **-66.4** | **0.0** | **0.60** |
| 1000 | 6 | Pasture / Mixed Forest / Semi-Natural/ Abandoned Extraction Open / Tallgrass Savannah / Total Pollinator Habitat | -63.1 | 3.3 | 0.11 |
| 1250 | 5 | Deciduous Forest / Semi-Natural/ Abandoned Extraction Open / Tallgrass Woodland / Total Pollinator Habitat | -64.9 | 1.5 | 0.28 |
| **1500** | **5** | **Deciduous Forest / Semi-Natural/ Built Up Area Pervious / Abandoned Extraction Open / Total Pollinator Habitat** | **-65.5** | **0.0** | **0.55** |
| 1750 | 6 | Deciduous Forest / Semi-Natural/ Built Up Area Pervious / Abandoned Extraction Open / Consistent Forage Crop / Total Pollinator Habitat | -64.3 | 1.2 | 0.30 |
| 2000 | 6 | Deciduous Forest / Semi-Natural/ Built Up Area Pervious / Abandoned Extraction Open / Consistent Forage Crop / Total Pollinator Habitat | -62.8 | 2.7 | 0.14 |
| **Solitary Ground Nesters Proportional Abundance** | | |  |  |  |
| 250 | 3 | Marsh / Tallgrass Woodland / Tallgrass Savannah | -93.8 | 31.4 | 0.00 |
| 300 | 3 | Marsh / Tallgrass Woodland / Tallgrass Savannah | -93.8 | 31.4 | 0.00 |
| 350 | 2 | Marsh / Tallgrass Woodland | -106.9 | 18.3 | 0.00 |
| 400 | 5 | Forest / Marsh / Built Up Area Pervious / Consistent Forage Crop / Tallgrass Woodland | -112.2 | 13.0 | 0.00 |
| 450 | 5 | Coniferous Forest / Thicket Swamp / Marsh / Tallgrass Woodland / Tallgrass Savannah | -117.2 | 8.0 | 0.02 |
| **500** | **5** | **Thicket Swamp / Marsh / Hedge Row / Semi-Natural/ Tallgrass Woodland** | **-125.2** | **0.0** | **1.00** |
| **750** | **5** | **Coniferous Forest / Marsh / Plantation / Hedge Row / Semi-Natural** | **-130.5** | **0.0** | **1.00** |
| 1000 | 4 | Coniferous Forest / Deciduous Forest / Marsh / Tallgrass Savannah | -129.8 | 0.7 | 0.72 |
| 1250 | 4 | Coniferous Forest / Deciduous Forest / Marsh / Tallgrass Woodland | -128.8 | 1.7 | 0.43 |
| 1500 | 3 | Marsh / Plantation / Tallgrass Woodland | -127.8 | 1.4 | 0.51 |
| **1750** | **3** | **Marsh / Plantation / Tallgrass Woodland** | **-129.1** | **0.0** | **1.00** |
| 2000 | 3 | Marsh / Plantation / Tallgrass Woodland | -128.9 | 0.2 | 0.90 |
| **Social Ground Nesters Proportional Abundance** | | |  |  |  |
| 250 | 6 | Treed Swamp / Pasture / Marsh / Plantation / Abandoned Extraction Open / Total Pollinator Habitat | -93.3 | 25.1 | 0.00 |
| 300 | 6 | Treed Swamp / Pasture / Marsh / Plantation / Abandoned Extraction Open / Total Pollinator Habitat | -95.3 | 23.1 | 0.00 |
| 350 | 6 | Treed Swamp / Forest / Mixed Forest / Marsh / Consistent Forage Crop / Tallgrass Savannah | -107.2 | 11.2 | 0.00 |
| 400 | 5 | Treed Swamp / Mixed Forest / Marsh / Consistent Forage Crop / Tallgrass Savannah | -110.6 | 7.8 | 0.02 |
| 450 | 6 | Forest / Thicket Swamp / Marsh / Hedge Row / Consistent Forage Crop / Tallgrass Woodland | -112.4 | 6.0 | 0.05 |
| **500** | **6** | **Treed Swamp / Coniferous Forest / Mixed Forest / Marsh / Built Up Area Pervious / Tallgrass Savannah** | **-118.4** | **0.0** | **1.00** |
| 750 | 5 | Treed Swamp / Pasture / Mixed Forest / Plantation / Tallgrass Savannah | -127.4 | 0.7 | 0.71 |
| 1000 | 5 | Coniferous Forest / Mixed Forest / Built Up Area Pervious / Tallgrass Savannah / Total Pollinator Habitat | -127.8 | 0.3 | 0.86 |
| **1250** | **6** | **Treed Swamp / Pasture / Coniferous Forest / Marsh / Semi-Natural/ Tallgrass Savannah** | **-128.0** | **0.0** | **1.00** |
| 1500 | 4 | Pasture / Marsh / Tallgrass Woodland / Tallgrass Savannah | -125.3 | 2.9 | 0.23 |
| 1750 | 4 | Pasture / Marsh / Tallgrass Woodland / Treed Sand Dune | -125.9 | 2.3 | 0.32 |
| **2000** | **6** | **Pasture / Coniferous Forest / Marsh / Hedge Row / Tallgrass Woodland / Treed Sand Dune** | **-128.2** | **0.0** | **1.00** |
| **Cavity Nesters Proportional Abundance** | | |  |  |  |
| 250 | 6 | Treed Swamp / Pasture / Forest / Marsh / Semi-Natural/ Consistent Foraging Crop | -117.4 | 29.7 | 0.00 |
| 300 | 6 | Treed Swamp / Forest / Marsh / Plantation / Semi-Natural/ Consistent Foraging Crop | -118.3 | 28.8 | 0.00 |
| 350 | 5 | Marsh / Semi-Natural/ Built Up Area Pervious / Consistent Foraging Crop / Total Pollinator Habitat | -120.4 | 26.8 | 0.00 |
| 400 | 6 | Treed Swamp / Mixed Forest / Thicket Swamp / Marsh / Semi-Natural/ Consistent Foraging Crop | -123.1 | 24.1 | 0.00 |
| 450 | 6 | Treed Swamp / Forest / Marsh / Semi-Natural/ Built Up Area Pervious / Consistent Foraging Crop | -139.0 | 8.2 | 0.02 |
| **500** | **6** | **Forest / Marsh / Hedge Row / Semi-Natural/ Consistent Foraging Crop / Total Pollinator Habitat** | **-147.2** | **0.0** | **1.00** |
| 750 | 6 | Coniferous Forest / Deciduous Forest / Marsh / Built Up Area Pervious / Consistent Foraging Crop / Total Pollinator Habitat | -146.6 | 1.8 | 0.41 |
| 1000 | 4 | Pasture / Coniferous Forest / Marsh / Built Up Area Pervious | -144.5 | 3.9 | 0.14 |
| **1250** | **6** | **Pasture / Forest / Coniferous Forest / Marsh / Abandoned Extraction Open / Tallgrass Savannah** | **-148.4** | **0.0** | **1.00** |
| **1500** | **6** | **Forest / Coniferous Forest / Mixed Forest / Marsh / Hedge Row / Total Pollinator Habitat** | **-156.6** | **0.0** | **1.00** |
| 1750 | 6 | Pasture / Forest / Coniferous Forest / Deciduous Forest / Marsh / Plantation | -154.2 | 2.4 | 0.30 |
| 2000 | 6 | Pasture / Forest / Coniferous Forest / Deciduous Forest / Marsh / Plantation | -151.9 | 4.7 | 0.09 |
| ***Bombus* Species Proportional Abundance** | | | |  |  |
| 250 | 2 | Deciduous Forest / Tallgrass Savannah | -109.6 | 66.3 | 0.00 |
| 300 | 4 | Coniferous Forest / Deciduous Forest / Abandoned Extraction Open / Tallgrass Savannah | -115.1 | 60.9 | 0.00 |
| 350 | 2 | Deciduous Forest / Tallgrass Savannah | -137.6 | 38.3 | 0.00 |
| 400 | 6 | Coniferous Forest / Marsh / Built Up Area Pervious / Abandoned Extraction Open / Tallgrass Woodland / Tallgrass Savannah | -197.3 | -21.3 | >1.0 |
| 450 | 6 | Forest / Deciduous Forest / Marsh / Built Up Area Pervious / Abandoned Extraction Open / Tallgrass Woodland | -149.9 | 26.1 | 0.00 |
| **500** | **6** | **Forest / Deciduous Forest / Marsh / Built Up Area Pervious / Consistent Forage Crop / Tallgrass Woodland** | **-176.0** | **0.0** | **1.00** |
| 750 | 5 | Coniferous Forest / Deciduous Forest / Marsh / Tallgrass Woodland / Total Pollinator Habitat | -185.9 | 26.6 | 0.00 |
| 1000 | 6 | Forest / Mixed Forest / Thicket Swamp / Marsh / Plantation / Abandoned Extraction Vegetated | -202.4 | 10.2 | 0.01 |
| **1250** | **6** | **Forest / Mixed Forest / Thicket Swamp / Marsh / Plantation / Abandoned Extraction Vegetated** | **-212.5** | **0.0** | **1.00** |
| **1500** | **3** | **Forest / Hedge Row / Abandoned Extraction Vegetated** | **-170.1** | **0.0** | **1.00** |
| 1750 | 6 | Forest / Deciduous Forest / Thicket Swamp / Marsh / Tallgrass Woodland / Treed Sand Dune | -156.9 | 13.2 | 0.00 |
| 2000 | 6 | Forest / Thicket Swamp / Marsh / Built Up Area Pervious / Tallgrass Woodland / Treed Sand Dune | -158.4 | 11.7 | 0.00 |
| **Cleptoparasite Proportional Abundance** | | | |  |  |
| 250 | 1 | Tallgrass Woodland | -135.3 | 39.9 | 0.00 |
| 300 | 1 | Tallgrass Woodland | -135.3 | 40.0 | 0.00 |
| 350 | 3 | Pasture / Consistent Forage Crop / Tallgrass Woodland | -155.9 | 19.4 | 0.00 |
| 400 | 1 | Tallgrass Woodland | -150.4 | 24.9 | 0.00 |
| 450 | 3 | Coniferous Forest / Thicket Swamp / Tallgrass Woodland | -161.1 | 14.1 | 0.00 |
| **500** | **2** | **Deciduous Forest / Tallgrass Woodland** | **-175.3** | **0.0** | **1.00** |
| 750 | 2 | Tallgrass Woodland / Tallgrass Savannah | -182.6 | 0.0 | 1.00 |
| 1000 | 2 | Tallgrass Woodland / Tallgrass Savannah | -182.6 | 0.0 | 1.00 |
| **1250** | **2** | **Tallgrass Woodland / Tallgrass Savannah** | **-182.6** | **0.0** | **1.00** |
| 1500 | 2 | Tallgrass Woodland / Tallgrass Savannah | -182.6 | 0.0 | 1.00 |
| 1750 | 2 | Tallgrass Woodland / Treed Sand Dune | -182.6 | 0.0 | 1.00 |
| **2000** | **2** | **Tallgrass Woodland / Treed Sand Dune** | **-182.6** | **0.0** | **1.00** |

Table S8. List of parameters of the best fit model at spatial distances <500m for species richness (top row) and abundance (bottom row) of each species groups: solitary ground nesters, social ground nesters, cavity nesters, *Bombus* spp. and cleptoparasites. The number of habitat parameters and name of habitat parameters used in each model are listed. Coefficients are based on log-transformed data and in bold where 95% CIs do not include 0. Negative coefficients suggest habitat type has a negative impact on expected species richness and proportional abundance of functional guild within specific spatial scale. Greyed out habitat types were not model parameters in any candidate models for any functional guild.

|  | Solitary ground | | |  | Social ground | | |  | Cavity nesters | | |  | *Bombus* spp. | | |  | Cleptoparasites | | | |
| --- | --- | --- | --- | --- | --- | --- | --- | --- | --- | --- | --- | --- | --- | --- | --- | --- | --- | --- | --- | --- |
| Land Classes | *β* | Lower CI | Upper CI |  | *β* | Lower CI | Upper CI |  | *β* | Lower CI | Upper CI |  | *β* | Lower CI | Upper CI |  | *β* | Lower CI | | Upper CI |
| Aband Extract Veg |  |  |  |  |  |  |  |  |  |  |  |  |  |  |  |  |  | |  |  |
| Aband Extract Open |  |  |  |  | **-0.35** | **-0.45** | **-0.26** |  | **-0.31** | **-0.40** | **-0.23** |  |  |  |  |  | **-0.66** | | **-0.70** | **-0.62** |
| Built Up Pervious |  |  |  |  |  |  |  |  |  |  |  |  |  |  |  |  |  | |  |  |
|  |  |  |  |  |  |  |  |  |  |  |  |  | **0.67** | **0.20** | **1.14** |  |  | |  |  |
| Coniferous Forest |  |  |  |  |  |  |  |  |  |  |  |  | **1.27** | **0.97** | **1.56** |  |  | |  |  |
| Consistent Forage Crop | **-0.12** | **-0.17** | **-0.07** |  | **-0.29** | **-0.49** | **-0.09** |  | **-0.36** | **-0.46** | **-0.25** |  |  |  |  |  | **-0.22** | | **-0.40** | **-0.05** |
|  |  |  |  |  | **0.36** | **0.26** | **0.47** |  | **-0.58** | **-0.70** | **-0.45** |  | 0.14 | -0.01 | 0.28 |  |  | |  |  |
| Deciduous Forest | **0.09** | **0.01** | **0.17** |  | **-1.04** | **-1.67** | **-0.41** |  | **0.41** | **0.28** | **0.54** |  |  |  |  |  |  | |  |  |
|  |  |  |  |  |  |  |  |  |  |  |  |  | **0.40** | **0.06** | **0.74** |  | **-0.29** | | **-0.46** | **-0.11** |
| Forest | 0.36 | -0.32 | 1.04 |  |  |  |  |  | **0.58** | **0.31** | **0.85** |  | **0.64** | **0.45** | **0.82** |  |  | |  |  |
|  |  |  |  |  | **-0.26** | **-0.37** | **-0.15** |  | **0.60** | **0.27** | **0.93** |  | -0.28 | -0.65 | 0.10 |  |  | |  |  |
| Hedge Row |  |  |  |  | -0.31 | -0.77 | 0.16 |  |  |  |  |  |  |  |  |  |  | |  |  |
|  | 0.43 | -0.03 | 0.88 |  |  |  |  |  | -0.32 | -0.64 | 0.0 |  |  |  |  |  |  | |  |  |
| Marsh |  |  |  |  | **-0.37** | **-0.49** | **-0.25** |  | -0.23 | -0.42 | -0.04 |  |  |  |  |  |  | |  |  |
|  | **-0.24** | **-0.44** | **-0.03** |  | **0.33** | **0.25** | **0.40** |  | **-0.58** | **-0.83** | **-0.33** |  | **0.47** | **0.16** | **0.77** |  |  | |  |  |
| Mixed Forest |  |  |  |  |  |  |  |  |  |  |  |  |  |  |  |  |  | |  |  |
|  |  |  |  |  | **-0.57** | **-0.74** | **-0.41** |  |  |  |  |  |  |  |  |  |  | |  |  |
| Pasture |  |  |  |  |  |  |  |  |  |  |  |  | **-0.47** | **-0.61** | **-0.32** |  | **-0.30** | | **-0.54** | **-0.06** |
| Plantation |  |  |  |  |  |  |  |  |  |  |  |  |  |  |  |  |  | |  |  |
| Semi-Natural |  |  |  |  |  |  |  |  |  |  |  |  |  |  |  |  | **-0.28** | | **-0.36** | **-0.20** |
|  | -0.29 | -0.60 | 0.02 |  |  |  |  |  | **0.50** | **0.30** | **0.70** |  |  |  |  |  |  | |  |  |
| Tallgrass Savannah | -0.93 | -1.90 | 0.04 |  | 1.08 | 0.597 | 1.570 |  |  |  |  |  |  |  |  |  |  | |  |  |
|  |  |  |  |  | **-0.52** | **-0.64** | **-0.40** |  |  |  |  |  |  |  |  |  |  | |  |  |
| Tallgrass Wood |  |  |  |  |  |  |  |  |  |  |  |  | **-0.92** | **-1.21** | **-0.63** |  |  | |  |  |
|  | **0.65** | **0.47** | **0.82** |  |  |  |  |  |  |  |  |  | **-0.51** | **-0.88** | **-0.15** |  | **0.87** | | **0.78** | **0.96** |
| Thicket Swamp | **-0.22** | **-0.39** | **-0.06** |  |  |  |  |  |  |  |  |  | **0.16** | **0.06** | **0.27** |  |  | |  |  |
|  | **0.34** | **0.05** | **0.63** |  |  |  |  |  |  |  |  |  |  |  |  |  |  | |  |  |
| Treed Sand |  |  |  |  |  |  |  |  |  |  |  |  |  |  |  |  |  | |  |  |
| Treed Swamp | **0.03** | **0.02** | **0.03** |  |  |  |  |  | **0.50** | **0.36** | **0.64** |  |  |  |  |  |  | |  |  |
| Total Pollinator Habitat |  |  |  |  |  |  |  |  |  |  |  |  | **-0.24** | **-0.41** | **-0.08** |  | **0.47** | | **0.27** | **0.66** |
|  |  |  |  |  |  |  |  |  | **-0.38** | **-0.65** | **-0.11** |  |  |  |  |  |  | |  |  |

Table S9. List of parameters of the best fit model at spatial 750-1250m for species richness (top row) and abundance (bottom row) of each species groups: solitary ground nesters, social ground nesters, cavity nesters, *Bombus* spp. and cleptoparasites. The number of habitat parameters and name of habitat parameters used in each model are listed. Coefficients are based on log-transformed data and in bold where 95% CIs do not include 0. Negative coefficients suggest habitat type has a negative impact on expect species richness and proportional abundance of functional guild within specific spatial scale. Greyed out habitat types were not model parameters in any candidate models for any functional guild.

|  | Solitary ground | | |  | Social ground | | |  | Cavity nesters | | |  | *Bombus* spp. | | |  | Cleptoparasites | | | |
| --- | --- | --- | --- | --- | --- | --- | --- | --- | --- | --- | --- | --- | --- | --- | --- | --- | --- | --- | --- | --- |
| Land Classes | *β* | Lower CI | Upper CI |  | *β* | Lower CI | Upper CI |  | *β* | Lower CI | Upper CI |  | *β* | Lower CI | Upper CI |  | *β* | Lower CI | | Upper CI |
| Aband Extract Veg |  |  |  |  | **-0.61** | **-0.75** | **-0.46** |  |  |  |  |  |  |  |  |  |  | |  |  |
|  |  |  |  |  |  |  |  |  |  |  |  |  | **1.37** | **1.09** | **1.65** |  |  | |  |  |
| Aband Extract Open |  |  |  |  |  |  |  |  |  |  |  |  |  |  |  |  | **-0.43** | | **-0.48** | **-0.38** |
| Built Up Pervious | **0.26** | **0.12** | **0.39** |  |  |  |  |  | **0.43** | **0.16** | **0.69** |  |  |  |  |  |  | |  |  |
| Coniferous Forest |  |  |  |  |  |  |  |  |  |  |  |  | **1.02** | **0.60** | **1.43** |  |  | |  |  |
|  | **0.81** | **0.61** | **1.00** |  | **-0.62** | **-0.82** | **-0.41** |  | **-0.50** | **-0.74** | **-0.26** |  |  |  |  |  |  | |  |  |
| Consistent Forage Crop |  |  |  |  | **0.32** | **0.09** | **0.55** |  |  |  |  |  |  |  |  |  |  | |  |  |
| Deciduous Forest | **0.57** | **0.28** | **0.85** |  |  |  |  |  | **0.47** | **0.34** | **0.60** |  | **0.69** | **0.31** | **1.08** |  |  | |  |  |
|  |  |  |  |  |  |  |  |  | **-0.72** | **-0.94** | **-0.51** |  |  |  |  |  |  | |  |  |
| Forest |  |  |  |  | **0.91** | **0.55** | **1.28** |  |  |  |  |  | **0.50** | **0.23** | **0.77** |  | **0.29** | | **0.08** | **0.50** |
|  |  |  |  |  |  |  |  |  | **0.86** | **0.48** | **1.24** |  |  |  |  |  |  | |  |  |
| Hedge Row |  |  |  |  |  |  |  |  |  |  |  |  |  |  |  |  | 0.22 | | 0.00 | 0.44 |
|  | 0.35 | -0.10 | 0.79 |  |  |  |  |  |  |  |  |  |  |  |  |  |  | |  |  |
| Marsh | **-0.34** | **-0.44** | **-0.23** |  |  |  |  |  | **-0.21** | **-0.39** | **-0.02** |  |  |  |  |  |  | |  |  |
|  | **-0.32** | **-0.45** | **-0.20** |  | **0.19** | **0.04** | **0.35** |  | **-0.28** | **-0.47** | **-0.08** |  | **-0.72** | **-1.25** | **-0.18** |  |  | |  |  |
| Mixed Forest |  |  |  |  |  |  |  |  |  |  |  |  |  |  |  |  |  | |  |  |
|  |  |  |  |  |  |  |  |  |  |  |  |  | **-0.57** | **-0.94** | **-0.21** |  |  | |  |  |
| Pasture |  |  |  |  |  |  |  |  |  |  |  |  |  |  |  |  |  | |  |  |
|  |  |  |  |  | **0.31** | **0.03** | **0.59** |  | **-0.30** | **-0.55** | **-0.05** |  | **-0.22** | **-0.35** | **-0.08** |  |  | |  |  |
| Plantation |  |  |  |  |  |  |  |  |  |  |  |  |  |  |  |  |  | |  |  |
|  | **-0.22** | **-0.36** | **-0.08** |  |  |  |  |  | **0.34** | **0.15** | **0.53** |  | **0.56** | **0.21** | **0.91** |  |  | |  |  |
| Semi-Natural | -0.12 | -0.27 | 0.04 |  |  |  |  |  | **-0.26** | **-0.44** | **-0.09** |  |  |  |  |  | **-0.31** | | **-0.51** | **-0.12** |
|  | -0.32 | -0.73 | 0.09 |  | **0.30** | **0.06** | **0.54** |  |  |  |  |  |  |  |  |  |  | |  |  |
| Tallgrass Savannah | **-0.33** | **-0.56** | **-0.09** |  |  |  |  |  |  |  |  |  |  |  |  |  |  | |  |  |
|  |  |  |  |  | **-0.26** | **-0.35** | **-0.17** |  |  |  |  |  |  |  |  |  | **1.86** | | **1.72** | **1.99** |
| Tallgrass Wood |  |  |  |  |  |  |  |  |  |  |  |  | **-0.56** | **-0.90** | **-0.21** |  |  | |  |  |
|  |  |  |  |  |  |  |  |  |  |  |  |  |  |  |  |  | **-1.41** | | **-1.49** | **-1.33** |
| Thicket Swamp |  |  |  |  | **-0.44** | **-0.54** | **-0.33** |  |  |  |  |  |  |  |  |  | **0.26** | | **0.12** | **0.41** |
|  |  |  |  |  |  |  |  |  |  |  |  |  | **0.79** | **0.30** | **1.28** |  |  | |  |  |
| Treed Sand |  |  |  |  |  |  |  |  |  |  |  |  |  |  |  |  |  | |  |  |
| Treed Swamp | **0.73** | **0.57** | **0.90** |  | **0.47** | **0.22** | **0.71** |  | **0.43** | **0.29** | **0.57** |  | **0.39** | **0.12** | **0.65** |  |  | |  |  |
|  |  |  |  |  | **0.62** | **0.12** | **1.11** |  |  |  |  |  |  |  |  |  |  | |  |  |
| Total Pollinator Habitat |  |  |  |  |  |  |  |  |  |  |  |  | **-1.22** | **-1.72** | **-0.72** |  | **0.59** | | **0.40** | **0.78** |

Table S10. List of parameters of the best fit model at spatial category >1500m for species richness (top row) and abundance (bottom row) of each species groups: solitary ground nesters, social ground nesters, cavity nesters, *Bombus* spp. and cleptoparasites. The number of habitat parameters and name of habitat parameters used in each model are listed. Coefficients are standardized and in bold where 95% CIs do not include 0. Negative coefficients suggest habitat type has a negative impact on expect species richness and proportional abundance of functional guild within specific spatial scale. Greyed out habitat types were not model parameters in any candidate models for any functional guild.

|  | Solitary ground | | |  | Social ground | | |  | Cavity nesters | | |  | *Bombus* spp. | | |  | Cleptoparasites | | | |
| --- | --- | --- | --- | --- | --- | --- | --- | --- | --- | --- | --- | --- | --- | --- | --- | --- | --- | --- | --- | --- |
| Land Classes | *β* | Lower CI | Upper CI |  | *β* | Lower CI | Upper CI |  | *β* | Lower CI | Upper CI |  | *β* | Lower CI | Upper CI |  | *β* | Lower CI | | Upper CI |
| Aban. Extract Veg |  |  |  |  |  |  |  |  |  |  |  |  | **-0.27** | **-0.43** | **-0.10** |  |  | |  |  |
|  |  |  |  |  |  |  |  |  |  |  |  |  | **0.81** | **0.31** | **1.31** |  |  | |  |  |
| Aban. Extract Open | **-0.51** | **-0.64** | **-0.38** |  |  |  |  |  |  |  |  |  |  |  |  |  | **-0.19** | | **-0.31** | **-0.07** |
| Built Up Pervious |  |  |  |  |  |  |  |  | **0.36** | **0.15** | **0.56** |  | **0.49** | **0.30** | **0.68** |  | **0.32** | | **0.12** | **0.53** |
| Coniferous Forest |  |  |  |  | **-0.60** | **-0.89** | **-0.30** |  |  |  |  |  | **0.61** | **0.22** | **0.99** |  |  | |  |  |
|  |  |  |  |  | 0.23 | -0.05 | 0.51 |  | **-1.21** | **-1.62** | **-0.80** |  |  |  |  |  |  | |  |  |
| Consistent Forage Crop |  |  |  |  |  |  |  |  |  |  |  |  |  |  |  |  | 0.18 | | -0.08 | 0.44 |
| Deciduous Forest | **-0.56** | **-1.03** | **-0.10** |  | **-0.80** | **-1.25** | **-0.34** |  | **-0.27** | **-0.44** | **-0.10** |  | **0.66** | **0.21** | **1.10** |  | 0.27 | | 0.00 | 0.53 |
| Forest | **1.04** | **0.69** | **1.40** |  | **0.78** | **0.34** | **1.21** |  | **0.64** | **0.44** | **0.84** |  |  |  |  |  |  | |  |  |
|  |  |  |  |  |  |  |  |  | **0.67** | **0.49** | **0.84** |  | -0.23 | -0.49 | 0.02 |  |  | |  |  |
| Hedge Row |  |  |  |  |  |  |  |  |  |  |  |  |  |  |  |  |  | |  |  |
|  |  |  |  |  | **0.27** | **0.03** | **0.52** |  | -0.28 | -0.56 | 0.00 |  | -0.31 | -0.68 | 0.07 |  |  | |  |  |
| Marsh |  |  |  |  |  |  |  |  |  |  |  |  |  |  |  |  |  | |  |  |
|  | **-0.38** | **-0.56** | **-0.19** |  | **0.47** | **0.27** | **0.67** |  | **-0.39** | **-0.52** | **-0.25** |  |  |  |  |  |  | |  |  |
| Mixed Forest |  |  |  |  |  |  |  |  |  |  |  |  |  |  |  |  |  | |  |  |
|  |  |  |  |  |  |  |  |  | **1.04** | **0.71** | **1.37** |  |  |  |  |  |  | |  |  |
| Pasture |  |  |  |  |  |  |  |  | **0.33** | **0.15** | **0.51** |  |  |  |  |  |  | |  |  |
|  |  |  |  |  | **0.35** | **0.02** | **0.69** |  |  |  |  |  |  |  |  |  |  | |  |  |
| Plantation | **2.09** | **1.32** | **2.86** |  |  |  |  |  |  |  |  |  |  |  |  |  |  | |  |  |
|  | **0.46** | **0.32** | **0.59** |  |  |  |  |  |  |  |  |  |  |  |  |  |  | |  |  |
| Semi-Natural |  |  |  |  |  |  |  |  | **-0.55** | **-0.70** | **-0.40** |  |  |  |  |  | **-0.52** | | **-0.65** | **-0.40** |
| Tallgrass Savannah |  |  |  |  |  |  |  |  |  |  |  |  |  |  |  |  |  | |  |  |
| Tallgrass Wood | **-9.16** | **-12.68** | **-5.65** |  | **-3.99** | **-4.96** | **-3.03** |  | **0.45** | **0.34** | **0.56** |  |  |  |  |  |  | |  |  |
|  | **0.48** | **0.41** | **0.54** |  |  |  |  |  |  |  |  |  |  |  |  |  | **3.809** | | **3.532** | **4.085** |
| Thicket Swamp |  |  |  |  | **-0.30** | **-0.41** | **-0.19** |  |  |  |  |  |  |  |  |  |  | |  |  |
| Treed Sand | **9.51** | **5.83** | **13.19** |  | **3.73** | **2.77** | **4.68** |  |  |  |  |  | **0.40** | **0.16** | **0.64** |  |  | |  |  |
|  |  |  |  |  |  |  |  |  |  |  |  |  |  |  |  |  | **-3.426** | | **-3.653** | **-3.198** |
| Treed Swamp |  |  |  |  |  |  |  |  |  |  |  |  |  |  |  |  |  | |  |  |
| Total Pollinator Habitat |  |  |  |  | **1.16** | **0.57** | **1.75** |  |  |  |  |  | **-1.20** | **-1.70** | **-0.71** |  | **0.41** | | **0.06** | **0.76** |
|  |  |  |  |  |  |  |  |  | **-0.49** | **-0.75** | **-0.24** |  |  |  |  |  |  | |  |  |

65.	OMNRF, Southern Ontario Land Resource Information System (SOLRIS) Version 2 .1, (Ontario Monistry of Natural Resources and Forestry Peterborough, Ontario, 2015).

66.	AAFC, Annual Crop Inventory Data, (Ottawa, Canada, 2016).

67.	ESRI, ArcGIS Desktop: Release 10.3.1, (Environmental System Research Institue, Redlands, CA., 2014).

68.	J. Hauke, T. Kossowski, Comparison of values of Pearson's and Spearman's correlation coefficients on the same sets of data. *Quaestiones Geographicae* **30**, 87-93 (2011).

69.	Addinsoft. (Boston, USA, 2020).

70.	R. C. Team, R Foundation for Statistical Computing; Vienna, Austria: 2014. *R: A language and environment for statistical computing*, 2013 (2018).

71.	S. Colla, E. Willis, L. Packer, Can green roofs provide habitat for urban bees (Hymenoptera: Apidae)? *Cities and the Environment* **2**, 1-12 (2009).

72.	H. Andrachuk, MSc, University of Waterloo (2014).

73.	J. C. Grixti, L. Packer, Changes in the bee fauna (Hymenoptera: Apoidea) of an old field site in southern Ontario, revisited after 34 years. *The* *Canadian Entomologist* **138**, 147-164 (2006).

74.	J. James, Native Bee Diversity in Conventional and Organic Hedgerows in Eastern

Ontario, MSc Thesis, University of Ottawa, Ottawa (2014).

75.	A. Pindar, The effect of fire disturbance on bee community composition in oak savannah habitat in southern Ontario, Canada, Ph.D. Thesis, York University, Toronto (2014).

76.	A. N. Taylor, P. M. Catling, Bees and butterflies in burned and unburned alvar woodland: evidence for the importance of postfire succession to insect pollinator diversity in an imperiled ecosystem. *The Canadian Field-Naturalist* **125**, 297-306 (2011).

77.	M. H. Richards *et al.*, Bee diversity in naturalizing patches of Carolinian grasslands in southern Ontario, Canada. *The Canadian Entomologist* **143**, 279-299 (2011).

78.	H. Lee, Southern Ontario Ecological Land Classification, (London, Ontario, 2008).
